# Spontaneous polyploids and antimutators compete during the evolution of mutator cells

**DOI:** 10.1101/718163

**Authors:** Maxwell A. Tracy, Mitchell B. Lee, Brady L. Hearn, Ian T. Dowsett, Luke C. Thurber, Jason Loo, Anisha M. Loeb, Kent Preston, Miles I. Tuncel, Niloufar Ghodsian, Anna Bode, Thao T. Tang, Andy R. Chia, Alan J. Herr

## Abstract

Heterozygous mutations affecting DNA polymerase (Pol) exonuclease domains and homozygous inactivation of mismatch repair (MMR) each generate “mutator” phenotypes capable of driving tumorigenesis. Cancers with both defects exhibit an explosive increase in mutation burden that appears to reach a threshold, consistent with selection acting against further mutation accumulation. In haploid yeast, simultaneous defects in polymerase proofreading and MMR select for “antimutator” mutants that suppress the mutator phenotype. We report here that spontaneous polyploids also escape this “error-induced extinction” and routinely out-compete antimutators in evolved haploid cultures. We performed similar experiments to explore how diploid yeast adapt to the mutator phenotype. We first evolved cells with homozygous mutations affecting proofreading and MMR, which we anticipated would favor tetraploid emergence. While tetraploids arose with a low frequency, in most cultures, a single antimutator clone rose to prominence carrying biallelic mutations affecting the polymerase mutator alleles. Variation in mutation rate between subclones from the same culture suggests there exists continued selection pressure for additional antimutator alleles. We then evolved diploid yeast modeling MMR-deficient cancers with the most common heterozygous exonuclease domain mutation (*POLE-P286R*). Although these cells grew robustly, within 120 generations, all subclones carried truncating or nonsynonymous mutations in the *POLE-P286R* homologous allele (*pol2-P301R*) that suppressed the mutator phenotype as much as 100-fold. Independent adaptive events in the same culture were common. Our findings suggest that analogous tumor cell populations may adapt to the threat of extinction by polyclonal mutations that neutralize the *POLE* mutator allele and preserve intra-tumoral genetic diversity for future adaptation.

## Introduction

The high fidelity of DNA replication and repair prevents mutations that might otherwise lead to cancer. As first proposed more than 40 years ago, cells with defects in these pathways exhibit an elevated mutation rate (“mutator phenotype”) and an accelerated path to malignancy (Loeb *et al.* 1974; Loeb 2016). The mutator hypothesis of cancer first gained support from the discovery that Lynch syndrome was caused by defects in mismatch repair (MMR), which normally corrects DNA replication errors (LYNCH *et al.* 2009). Mouse studies later demonstrated that loss of polymerase proofreading by the major leading and lagging strand DNA polymerases (Polε and Polδ, respectively) also promoted tumorigenesis (Goldsby *et al.* 2001; Albertson *et al.* 2009). More recently, cancer genome sequencing revealed exonuclease-domain mutations in the catalytic subunit genes for Polε (*POLE*) and Polδ (*POLD1*) in a subset of highly mutated colorectal and endometrial human cancers (Yoshida *et al.* 2011; Network 2012; Palles *et al.* 2012; Church *et al.* 2013; Kandoth *et al.* 2013). Heterozygous mutations in *POLE* predominate in these “ultra-hypermutated” cancers, with the most common *POLE* variant encoding a P286R substitution (Barbari *et al.* 2018). *POLE-P286R* confers strong mutator and cancer phenotypes when engineered into mice in the heterozygous state (Li *et al.* 2018). The corresponding allele in budding yeast (*pol2-P301R*) was found to confer a strong dominant mutator phenotype that far exceeds that of an exonuclease-null allele (*pol2-4*) (Kane and Shcherbakova 2014). Recent biochemical and crystallography experiments indicate that the yeast Polε P301R variant retains residual exonuclease activity but encodes a hyperactive polymerase that excels at mispair extension (Xing *et al.* 2019) due to occlusion of the DNA binding site of the exonuclease domain by the arginine substitution (Parkash *et al.* 2019). Many of the other *POLE* exonuclease domain alleles, which confer weaker mutator phenotypes than *pol2-P301R* when modeled in yeast, may also promote mispair extension rather than inhibit proof-reading (Barbari *et al.* 2018).

Tandem defects in proofreading and MMR pathways produce greater than additive increases in mutation rate in bacteria and yeast (Morrison *et al.* 1993; Schaaper 1993; Morrison and Sugino 1994) indicating that polymerase fidelity and mismatch repair act redundantly to limit replication errors. Correspondingly, some human tumors with heterozygous *POLE* or *POLD1* mutations also carry mutations affecting MMR (Network 2012; Kandoth *et al.* 2013). Far from incidental, the MMR defects appear to synergize with the polymerase variants to accelerate mutagenesis. In so doing, they create mutational signatures consistent with the interaction between these two pathways (Campbell *et al.* 2017; Haradhvala *et al.* 2018; Hodel *et al.* 2018). Likewise, patients with inherited biallelic MMR deficiency frequently develop rapidly mutating tumors with heterozygous mutations affecting the fidelity of Polε or Polδ (Shlien *et al.* 2015). Thus, powerful mutator phenotypes based on the synergistic interactions between polymerase fidelity and MMR defects represent a reoccurring event in cancer evolution.

As an evolutionary strategy, cells that employ a high mutation rate run the risk of error-induced extinction (*EEX*), where every one of their descendants eventually acquires a lethal mutation (Morrison *et al.* 1993; Fijalkowska and Schaaper 1996; Herr *et al.* 2011a). As an extreme manifestation of *EEX*, double mutant haploid yeast spores deficient in Polδ proofreading and MMR cease dividing within 10 cellular generations (Morrison *et al.* 1993). Strong mutator phenotypes can also drive *EEX* of diploid yeast, although the maximum tolerated mutation rate (error threshold) is higher than in haploids, consistent with genetic buffering by the diploid genome (Herr *et al.* 2014). Colony formation slows and becomes irregular once mutation rates are within an order of magnitude of the diploid error threshold. High mutation loads may inherently restrict cellular longevity. Recent lifespan experiments in yeast demonstrate that strong diploid mutators exhibit a form of genetic anticipation: the longer a lineage exists, the higher the mutation burden, and the shorter the cellular lifespan (Lee *et al.* 2019). Evidence suggests that error thresholds also exist in mammals. Mouse embryos completely deficient in both proofreading and MMR initiate development but fail in the second week of gestation (Albertson *et al.* 2009). Moreover, tumors with combined deficiencies in polymerase fidelity and MMR appear to reach an upper limit of ∼ 250 mutations/Mb, which has been interpreted as evidence for a mutation threshold in cancer (Shlien *et al.* 2015).

It is unclear whether strong mutator-driven cancers adapt to the threat of extinction and, if so, how. In yeast, one third of haploid *“eex”* mutants with combined defects in proofreading and MMR acquire intragenic “antimutator” mutations that suppress the mutator polymerase (Herr *et al.* 2011a; Williams *et al.* 2013). Polymerase antimutators also arise in bacteria and bacteriophage expressing strong mutator polymerases, suggesting that these adaptive mechanisms are highly conserved (Reha-Krantz 1988; Fijalkowska and Schaaper 1995). The yeast antimutator mutations encode amino acid substitutions in a wide variety of positions within the polymerase including the polymerase active site, DNA binding domains, polymerase structural elements, or even the exonuclease domain itself (Herr *et al.* 2011a; Herr *et al.* 2011b; Williams *et al.* 2013; Dennis *et al.* 2017). Since Polδ and Polε both play essential roles in DNA replication, these antimutator alleles found in haploid cells likely produce functional polymerases. Some substitutions may increase nucleotide selectivity while others may promote polymerase dissociation from mispaired primer termini, allowing extrinsic proofreading (Herr *et al.* 2011b). Evidence for extrinsic proofreading in yeast between Polδ and Polε exists in the literature (Morrison and Sugino 1994; Flood *et al.* 2015; Bulock *et al.* 2020) and processing of mispaired primer termini by other DNA repair pathways may also be possible (Herr *et al.* 2011b).

The majority of *eex* mutants in the above haploid yeast genetic screens, which utilized plasmid shuffling, carried genetic changes that were extragenic to the plasmid-borne mutator polymerase gene and were never mapped (Herr *et al.* 2011a; Williams *et al.* 2013). Here, we performed new genetic screens that would allow us to more easily identify these loci using Mendelian segregation of mutator and antimutator phenotypes. In the process, we discovered that many *eex* mutants were, in fact, not antimutators, but spontaneous polyploids, which are buffered from mutation accumulation as described above. This finding led us to explore the competition between antimutators and spontaneous polyploids during the evolution of diploid mutators. These diploid mutator strains did not display an initial growth defect but became subject to selection during propagation. This scenario models a population of mutator tumor cells having to adapt to the threat of extinction during cancer progression.

## Results

### Discovery of spontaneous polyploid *eex* mutants

The genetic screens that led to the discovery of spontaneous polyploid *eex* mutants began with diploid parent strains that were heterozygous for deletion of *MSH6 (msh6Δ)* and *pol3-01* (Polδ proofreading defective), or *msh2Δ* and *pol2-4* (Polε proofreading defective) (Figure S1A,B). After inducing the diploids to undergo sporulation (meiosis) we isolated *eex* mutants that suppressed the *EEX* phenotypes of freshly isolated double-mutant haploid cells. Both parental strains were heterozygous for two distinct transgene insertions at the *CAN1* locus: each *eex* mutant carried either *CAN1::natMX*, which confers sensitivity to canavanine and resistance to nourseothricin (NTC), or *can1Δ::HIS3*, which confers canavanine resistance and histidine prototrophy. We crossed all *eex* mutants to WT haploids with the opposite *CAN1* allele and mating type and then isolated diploid mapping strains by plating for His^+^ and NTC-resistant colonies. We sporulated the resulting mapping strains and although they routinely formed tetrads, the majority of mapping strains from both genetic screens failed to produce any viable spores (Figure S1B). Surprisingly, several mapping strains with intermediate spore viability produced progeny that carried both *CAN1* alleles. Flow cytometry revealed that these un-usual mapping strains were in fact tetraploids (4n) (Figure S1*B*), while the mapping strains with no viable spores were triploids (3n). We inferred that the original *eex* mutants for the 3n mapping strains were *MAT***a***/MAT***a** or *MAT*α/*MAT*α diploids and that the 4n strains were derived from *MAT***a**/*MAT***a**/*MAT***a** or *MAT*α/*MAT*α/*MAT*α triploid *eex* mutants. Since polyploid *eex* mutants arose in both genetic screens, this escape mechanism is a general one, likely related to the ability of polyploids to better withstand mutation accumulation.

### Evolution of haploid mutator cells

A tendency to pick larger candidate *eex* colonies could explain why polyploids were more common than antimutators in our genetic screens. To determine the relative success of these two adaptations in an unbiased manner, we isolated 89 independent *pol2-4 msh2Δ* spore clones by dissecting tetrads from *POL2/pol2-4 MSH2/msh2Δ* diploids on growth media that selected for the transgenes linked to the mutator alleles (Figure 1A). Freshly dissected *pol2-4 msh2Δ* haploids form colonies with an uneven perimeter, consistent with the onset of *EEX* and the emergence of *eex* mutant subclones with improved survival. After two days, we isolated four cells from the perimeter of each colony and moved them elsewhere on the plate to assess their colony forming capacity, mutation rate and ploidy (Figure S2A). Two-thirds of the cells isolated from the perimeters of the initial colonies failed to form a second colony — a manifestation of *EEX*. The 120 colonies from viable cells contained 12 that were uniformly polyploid (including one triploid (A4.1)) (Figure S2B, Dataset S1). To test for antimutators, we measured mutation rates of the haploid subclones that carried the *CAN1::natMX* allele. This involved subcloning these cells a second time to obtain replica colonies, which we assessed for ploidy. A high frequency of polyploid replica colonies prevented measurements for 12 sub-clones. Thirteen subclones exhibited the expected mutation rate for *pol2-4 msh2Δ* cells, while twelve subclones displayed an antimutator phenotype (Figure S2C). These results indicate that antimutators and polyploids emerge with similar frequencies during the initial growth of *pol2-4 msh2Δ* haploids and that polyploids continue to arise.

**Figure 1.**
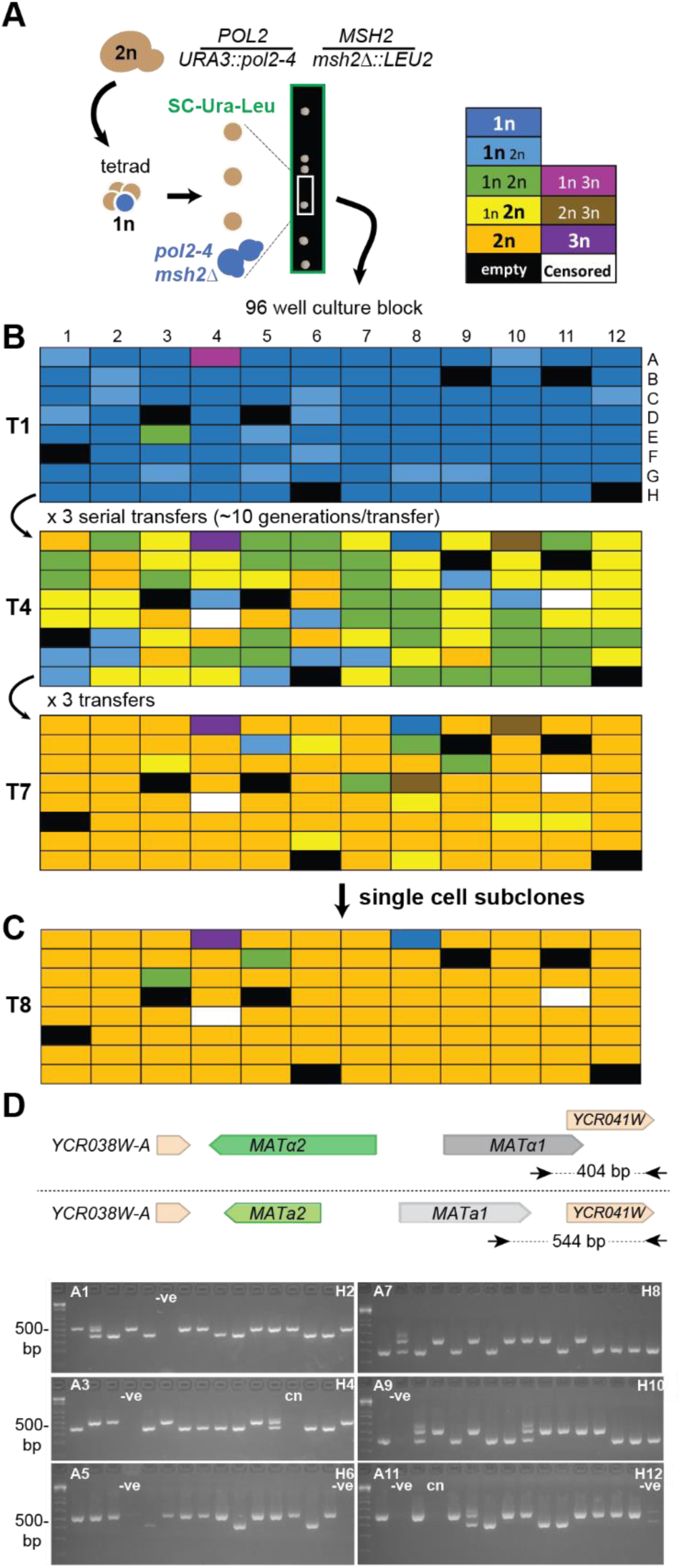
Spontaneous polyploids routinely emerge during the evolution of *pol2-4 msh2Δ* haploids. A) Isolation of *pol2-4 msh2Δ* haploids from sporulated parental diploid cells (2n). Diploid genotype indicated to the right. Arrows, from left to right, indicate manipulations of cells: sporulation, tetrad dissection, and inoculation into 96-well culture blocks. Following tetrad dissection, double mutant *pol2-4 msh2Δ* spores (blue cells) selectively form colonies on media lacking uracil and leucine (green box). B) Evolution of *pol2-4 msh2Δ* mutators. Independent isolates were cultured in 96-well format by diluting 1:500 with each transfer (T). Ploidy was assessed by flow cytometry at T1, T4, and T7 (see Dataset S1 for histograms). Color of boxes indicate the ploidy or mixture of ploidies of each culture (see top for key). C) Ploidy of single cell subclones isolated from T7. D) Mating-type of single cell subclones. *Top*, Schematic of *MAT***a** and *MAT*α mating type loci, PCR primers (arrows), and expected DNA sizes. *Bottom*, PCR products run on 2% agarose gels, labeled by their coordinates in the 96-well block; *-ve*, negative (no cells) controls; *cn*, culture censored during evolution. The 500bp fragment of the Invitrogen 1kb plus ladder is indicated to the left.

To investigate which *eex* strategy prevails during long-term growth, after removing individual cells for the above analysis, we suspended the original colonies in liquid growth media and evolved them in 96-well format with interspersed empty wells to serve as controls for contamination. We repeatedly sub-cultured the cells by diluting them 1:500 (∼10^5^ cells/inoculum) and growing them to saturation (∼10^8^ cells), yielding an estimated 10-12 generations per transfer. We assessed ploidy at the initial time point (T1) and then after the 4th (T4) and the 7th transfer (T7). The ploidies of the initial colony suspensions were overwhelming haploid, although 15 out of 89 cultures at this early stage showed evidence of a minor diploid subclone (Figure 1B, Dataset S1). The colony suspension from the above triploid subclone (A4) contained both haploid and triploid cells in equal proportions. Prolonged growth of the original cultures revealed profound fitness differences between haploid antimutators and spontaneous polyploids (Figure 1B). At T7 only 2 cultures remained haploid; 10 were mixtures of 1n and 2n cells, and 73 cultures were uniformly 2n or 3n. Thus, polyploids routinely win out over antimutators during the adaptation of haploid cells to the mutator phenotype.

### Mechanism of polyploidization in haploid mutator cultures

In the haploid evolution experiment, cells could have become polyploid by either mating or failed cytokinesis (Ganem *et al.* 2007). Although deletion of *Homing Endonuclease* (*HO*) stabilizes mating type in our studies, mating-type switching may still occur through random double-stranded DNA breaks at the *MAT* locus, which are then repaired using the transcriptionally silent *HML* (MATα) or *HMR* (MAT**a**) loci (Haber 2012). Conceivably, the rate of these events could be elevated in mutator cells. To determine the relative contributions of mating and spontaneous polyploidization, we isolated a single colony from each evolved culture and performed flow cytometry. Of the 87 isolates, there were 83 diploids, 1 haploid, 1 triploid, and 2 mixed cultures with both 1n and 2n cells (Figure 1C). Mating-type PCR assays revealed 41 *MAT***a***/MAT***a**, 36 *MAT*α/*MAT*α and 6 *MAT***a***/MAT*α diploids (Figure 1D). Three of the *MAT***a**/*MAT*α isolates were heterozygous at the *CAN1* locus (*CAN1::natMX/can1Δ::HIS3*) (Dataset S2), providing evidence for rare mating events between cells from neighboring wells. The three remaining *MAT***a***/MAT*α cells were homozygous for *CAN1::natMX* or *can1Δ::HIS3* and could have arisen through contamination or mating-type switching. We specifically investigated the frequency of mating-type switching with both *MAT***a** and *MAT*α cells. Diploids formed in both same-sex pairings with similar frequencies. But while *MAT***a** *x MAT***a** crosses nearly always gave rise to *MAT***a***/MAT*α diploids, *MAT*α *x MAT*α crosses produced *MAT*α*/MAT*α diploids (Figures S3, S4). Same-sex mating has previously been observed (Strathern *et al.* 1981). Given the similarity in frequencies, a dsDNA break at the *MAT* locus likely initiates both mating-type switching of *MAT***a** cells and *MAT*α same-sex mating (see Methods). The 3 *MAT***a***/MAT*α clones that were homozygous for the *CAN1::natMX* or *can1Δ::HIS3* allele, which arose by mating type switching, suggests that only a small percentage of the 36 *MAT*α*/MAT*α cultures in the evolution experiment arose by same-sex mating. Together these observations argue that the majority of *MAT***a***/MAT***a** or *MAT*α/*MAT*α isolates in our evolving populations arose by spontaneous polyploidization due to failed cytokinesis, which recent studies indicate occurs with a frequency of 7 × 10^−5^ in haploid yeast (Harari *et al.* 2018). Since tetraploids have been observed in various cancers, we reasoned that spontaneous polyploidization could conceivably represent a conserved mechanism of escape from *EEX* for mutator-driven tumors.

### Tetraploidization further protects spontaneous diploids

As a first test of whether tetraploidy confers an advantage in a diploid mutator population we utilized the above spontaneous *pol2-4/pol2-4 msh2Δ/msh2Δ* diploids. We isolated *MAT***a***/MAT***a** and *MAT*α/*MAT*α cells from different evolved cultures by microdissection and mated their daughter cells. We moved the parental diploid cells as well as the resulting tetraploid zygotes to distinct locations on the agar plates to form colonies (Figure 2A). We then mixed each parent diploid strain separately with their tetraploid progeny at different ratios (100:1, 10:1, 1:1, 1:10, 1:100) and grew them in competition. The tetraploids overtook the parental mutator diploids in 7 out of 8 competitions. These experiments suggest that tetraploid mutators would potentially have a sizeable fitness advantage over the diploid mutator population from which they arise. An important caveat to this experiment, however, is that both diploid parents likely carry homozygous mutations that first arose in the original haploid mutators. These genetic liabilities would became heterozygous in their tetraploid offspring. In diploid cells that spontaneously acquire a mutator phenotype, mutations would accumulate from the beginning in a buffered, heterozygous state.

**Figure 2.**
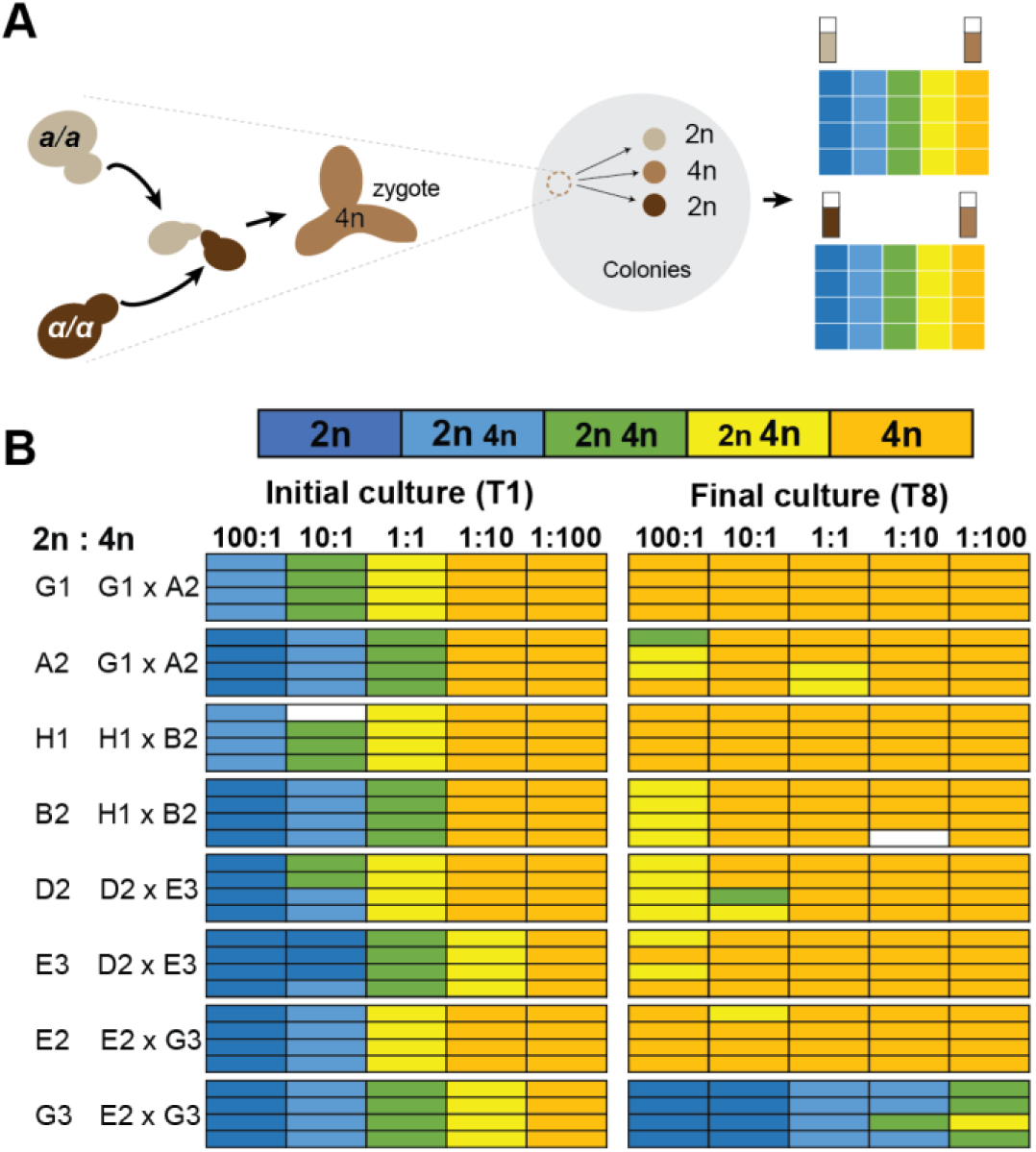
Competition between diploid mutator parents and tetraploid offspring. A) Experimental design. *MAT*α/*MAT*α and *MAT***a**/*MAT***a** subclones from the *pol2-4 msh2Δ* evolved cultures (see Figure 1B) were mated. Cell suspensions of the resulting tetraploid and parental diploid colonies were counted, mixed and then competed through 8 serial transfers. B) Ploidy measurements after the first (T1) and eighth (T8) serial transfers (see Dataset 1 for histograms). Letter and numbers to the left indicate location in the 96 well culture block described in Figure.1. Color of boxes indicate the ploidy or mixture of ploidies of each culture (see top).

### Adaptation of diploid yeast to a strong sublethal mutator phenotype

To explore how diploid cells with a near WT mutation burden adapt to a strong mutator phenotype, we began with *pol3-01/pol3-01 msh6Δ/msh6Δ* cells, which form robust colonies despite having a mutation rate (1 × 10^−3^ Can^R^ mutants/division) just below the diploid error threshold (Figure 3A)(Herr *et al.* 2014). Using a mathematical model in which the decrease in fitness was driven solely by lethal homozygous inactivation of essential genes (Herr *et al.* 2014), we estimated that a tetraploid or antimutator *eex* mutant would overtake a *pol3-01/pol3-01 msh6Δ/msh6Δ* diploid culture in 200 generations (Figure S5, Dataset 1). With two sets of homozygous mutator alleles driving the mutator phenotype, we anticipated tetraploids would be at their greatest advantage relative to antimutators.

**Figure 3.**
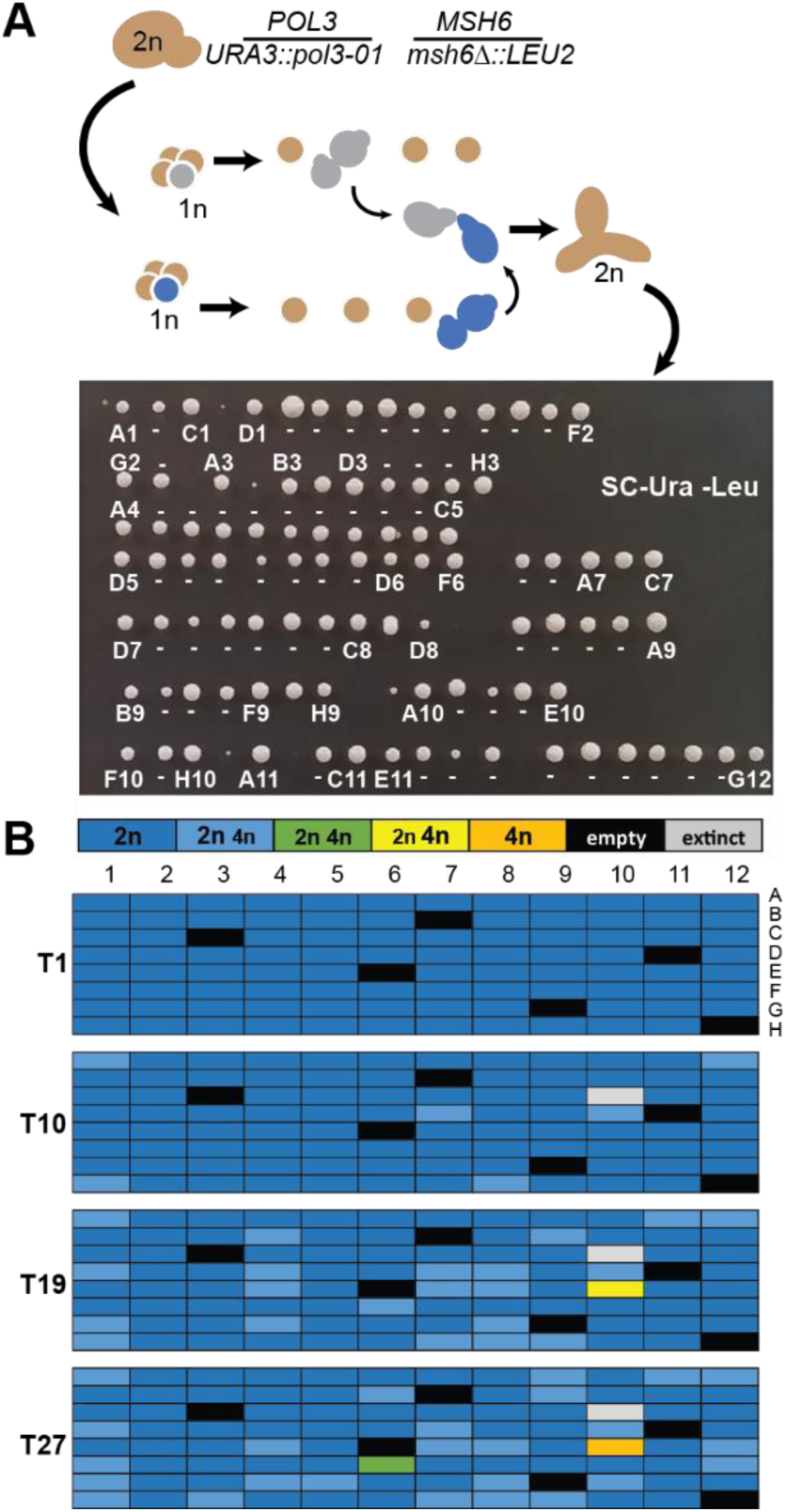
Strong diploid mutators maintain diploidy during evolution. A) Isolation of *pol3-01/pol3-01 msh6Δ/msh6Δ* zygotes. *Top*, Double mutant *pol3-01 msh6Δ* haploids, identified by dissecting tetrads on selective media (Sc-Ura-Leu), were mated to form zygotes. Zygotes were then moved to new locations on the plate to form colonies. *Bottom*, Growth phenotypes of the initial zygote colonies. Colonies were picked from left to right. Those used for evolution are indicated by either their 96-well coordinate or a dash (-). Colonies without a label or dash were not evolved. B) Ploidy of evolved cultures after one (T1), ten (T10), nineteen (T19), and twenty-seven (T27) serial transfers (see Dataset S1 for histograms). Color of boxes indicate the ploidy or mixture of ploidies of each culture as depicted in the key at the top.

To set up the evolution experiment, we first isolated numerous independent *pol3-01/pol3-01 msh6Δ/msh6Δ* zygotes by mating *pol3-01 msh6Δ* haploids that were freshly dissected from the *pol3-01/POL3 msh6Δ/MSH6* parental strain (Figure 3A). We evolved 89 zygote clones through ∼300 generations, monitoring ploidy at the T1, T10, T19, and T27 transfers (Figure 3B). A single pure tetraploid culture emerged at the end of the experiment. The remaining cultures were overwhelmingly diploid, suggesting that they adapted through antimutator mutations. To assay for antimutator phenotypes, we isolated independent subclones from T25 *CAN1::natMX/can1Δ::HIS3* cultures, assessed their ploidy, and measured mutation rates. Almost all subclones were diploids (Dataset 1) and exhibited a clear antimutator phenotype (Figure 4A). Antimutators also dominated the T13 cultures and were evident as early as T1 (Figure 4A). T1 cells from every culture formed variably sized colonies, which were notably smaller than T13 colonies, suggesting that strong selection for *eex* mutants exists by ∼30 cellular generations (20 generations/colony and 10 generations for T1) (Figure S6).

**Figure 4.**
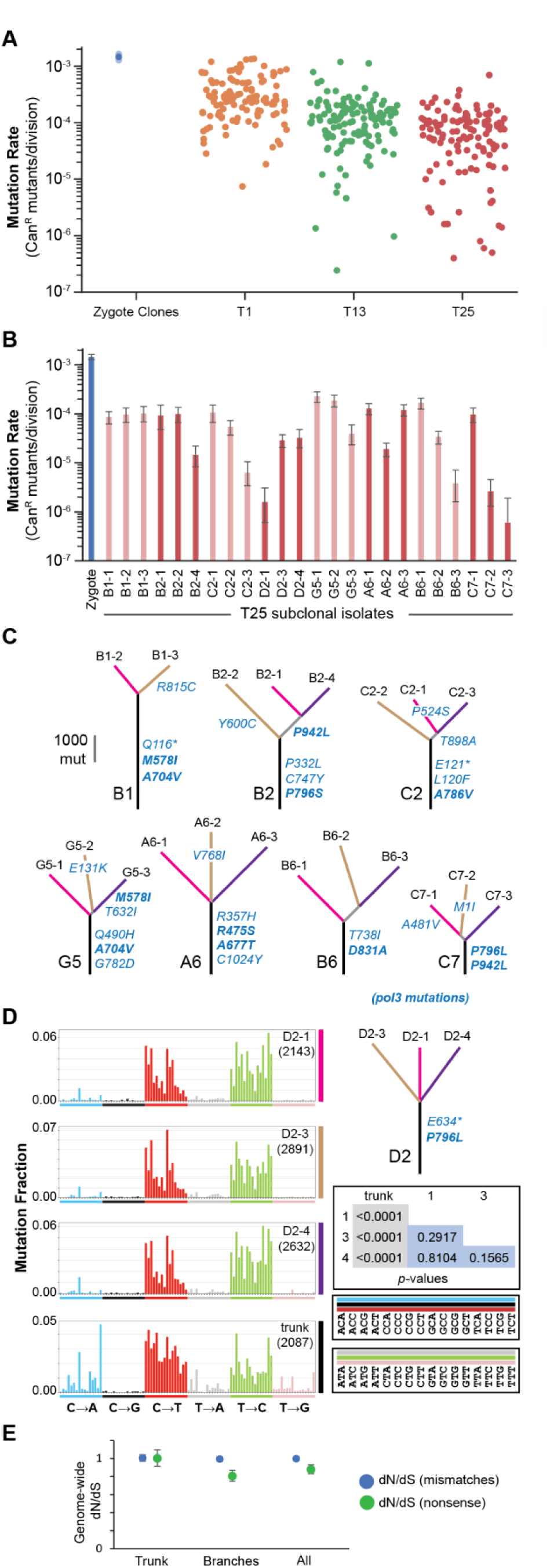
Strong diploid mutators adapt through antimutator alleles. A) Mutation rates of *pol3-01/pol3-01 msh6Δ/msh6Δ* subclones. Single cell subclones were isolated after the first (T1), thirteenth (T13), and twenty-fifth (T25) transfers. Mutation rates were determined by a canavanine resistance (Can^R^) fluctuation assay (see Dataset S1 for mutation rates and 95% confidence intervals). Solid circles, mutation rates; transparent circles, 95% confidence intervals. B) Variation in mutation rates depicted in (A) of T25 subclones from the same culture. Alternating shades of red indicate groups of subclones. C) Phylogenetic trees of subclones depicted in (B) based on whole genome sequencing. Observed *pol3* mutations are in blue lettering using single letter amino acid codes: *, nonsense mutation; bold type, known or likely antimutators). D) Representative mutation spectra of shared (trunk) and unique mutations (see Figure S7 for mutation spectra of all trees in (C)). Each mutation site (represented here by the pyrimidine base T or C) was categorized into one of the 96 mutation types created by subdividing the 6 general mutation types (C**→**A, C**→**G, C**→**T, T**→**A, T**→**C, T**→**G) into the 16 possible tri-nucleotide contexts (see boxes to the lower right for contexts; blue, black, and red bars correspond to mutation sites involving C; gray, green and pink bars correspond to mutation sites involving T). Bar height corresponds to the fraction each mutation type represents of the total mutations (given in the right-hand corner of each plot). Inset table: *p*-values of the hypergeometric test of similarity between mutation spectra. E) Genome-wide (global) dN/dS ratios of trunk and branch mutations.

Independent T25 isolates from the same *pol3-01/pol3-01 msh6Δ/msh6Δ* culture sometimes displayed different mutation rates (Figure 4B). We wondered whether they represented independent adaptations, and therefore sequenced the genomes of eight sets of subclones. In each set, the subclones could be arranged into a phylogenetic tree with ∼1300 to 4500 shared single nucleotide variants (SNVs) in the trunk and ∼1400 to 3000 unique SNVs in the branches (Figure 4C,D). This indicates that a single adapted clone rose to prominence in each culture and then diverged. Thus, differences in mutation rates between subclones likely reflect the presence of additional antimutator mutations in some isolates but not others.

The mutations that delineate the phylogenetic trees bear the imprint of the mutator and antimutator phenotypes involved in the evolution of these cultures. We classified the shared and unique SNVs according to the 96 trinucleotide contexts defined by the six different mutation subtypes (C:G→A:T, C:G→G:C, C:G→T:A, T:A→A:T, T:A→C:G, T:A→G:C), flanked by all possible 5′ and 3′ nucleotides (Figure 4D). We then compared mutation spectra using the hypergeometric and cosine similarity tests. By both tests, the spectra of the trunk mutations were largely the same between different cultures, consistent with each culture beginning with a common mutator phenotype (Figures 4D, S7, Dataset S2). The spectrum of unique mutations in each branch of a given tree differed substantially from that of the trunk mutations (*p* < 0.0001), although generally not from each other, consistent with the antimutator phenotype arising prior to their divergence (Figure 4D, S7). The antimutator alleles caused particularly sharp decreases in the relative abundance of C:G→A:T mutations in the branches of all trees. Despite these similarities, the spectra of unique mutations from unrelated evolved cultures often differed statistically from each other, suggesting that the antimutator alleles altered replication fidelity in distinct ways (Figure 4D, Dataset S2). Thus, acquisition of an antimutator phenotype that fundamentally changes the mutation spectrum coincides with the key adaptive event in each culture.

To understand how the cells attenuated the mutator phenotype, we annotated the mutations (Dataset S2) and compared the affected genes. All 8 strain sets acquired multiple *pol3* mutations prior to their divergence (Figure 4C,D, see trunks). Polδ plays an essential role in DNA replication. Both *pol3-01* alleles contribute to the onset of error-induced extinction, but only one allele needs to remain functional for cells to survive. Thus, the premature nonsense codons observed in three trees (B1, C2, D2) contribute to mutator suppression by inactivating one of the two *pol3-01* alleles, which allows the other *pol3-01* allele with an antimutator mutation to serve as the sole source for Polδ. Many missense mutations affect the same amino acid residue as (or are identical to) known antimutator mutations (*A704V(2x), A786V, A677T, D831A, R475S, R815C*) (Herr *et al.* 2011a; Herr *et al.* 2014; Dennis *et al.* 2017). The other missense mutations affecting *pol3-01* may inactivate polymerase function or may be *bona fide* antimutator alleles that have not previously been identified. Candidate antimutator mutations include *P942L* and *M578I*, which were each observed in two independent lineages, as well as mutations affecting P796, which were isolated three independent times (*P796L*(2x) and *P796S*) — once in combination with a stop codon. In some lineages, additional *pol3* alleles were observed in the branches. In two cases (G5-3, A6-2), these may account for the variation in mutation rates between subclones from the same culture (Figure 4B), but in other cases they clearly do not (B2-4, C2-3, D2-1, B6-2, B6-3, C7-3), implicating other loci in the attenuation of mutation rates.

We looked for genes under positive selection by comparing the ratio of nonsynonymous and synonymous mutations (dN/dS) across the genome using *dNdscv*, which normalizes the observed dN/dS ratio by the 192-trinucleotide mutation spectra of each gene in order to assess whether the observed ratio departs from neutrality (dN/dS = 1) (Martincorena *et al.* 2017). We analyzed the mutations from the trunks of the 8 lineages and found that only *pol3* mutations are under positive selection (q = 0.004, corrected *p-*value using Benjamini and Hochberg’s false discovery rate)(Dataset S2). Mutations in other genes, within the resolution of this dataset, accumulate in a neutral fashion. Repeating this analysis for the 23 unique sets of mutations from the branches failed to reveal any other genes under strong selection despite evidence for their existence, suggesting that more observations are needed for sufficient statistical power, especially if there are multiple antimutator genes. Interestingly, when branch and trunk mutations were analyzed together, *pol3* mutations do not emerge from the multiple testing correction as significant, underscoring the importance of using phylogenetic information for the identification of candidate antimutators by dN/dScv.

Being able to classify mutations according to the phylogenetic tree also provided evidence for negative selection acting against the mutator phenotype. For subclasses of trunk mutations involving missense or nonsense mutations, which arose early during mutator evolution, the genome-wide (or global) dN/dS ratio was 1.0, indicative of neutrality (Figure 4E)(Dataset S2). However, the genome-wide dN/dS ratio in the branches for nonsense mutations was 0.8, indicating that negative selection begins to act against the most deleterious class of mutations during extended propagation of these lines.

### Modeling adaptation to the *POLE-P286R* mutator phenotype in cancer

Having established that diploid mutators rapidly evolve to avoid extinction in long-term culture, we sought to understand how cells would adapt to the most common *POLE* mutator allele in cancer, *P286R*. Most *POLE-P286R* tumors appear microsatellite-stable and bear mutational signatures that suggest that they are MMR proficient (Haradhvala *et al.* 2018). However, one *POLE-P286R* patient curated on cBioPortal (Cerami *et al.* 2012) carried two distinct null alleles of *MSH6* (TCGA-IB-7651)(Dataset 1) and a clear mutation signature consistent with synergy between *POLE-P286R* and MMR deficiency (Haradhvala *et al.* 2018). To model this extreme scenario, we took a similar approach as before and isolated numerous independent *pol2-P301R/POL2 msh2Δ/msh2Δ* zygotes by crossing freshly dissected *pol2-P301R msh2Δ* and *POL2 msh2Δ* haploid spores (Figure 5A). Whereas concurrently isolated double homozygous mutants formed small colonies of around 1 x 10^5^ cells, the *pol2-P301R/POL2 msh2Δ/msh2Δ* zygotes grew robustly (Figure 5B, Figure S8) and we evolved 81 independent clones through 10 transfers (∼120 generations). By T10, one culture had evolved a prominent tetraploid subclone (Figure 5C). To test for antimutator phenotypes in the others, we isolated three subclones from 44 *CAN1::natMX/can1Δ::HIS3* cultures. Genotyping revealed 1 out of the 132 isolates had lost the heterozygous *pol2-P301R* allele through mitotic recombination (H9-1). Most of the other isolates had mutation rates more than 10-fold lower than the original zygote clones, and many as low as the mitotic recombinant, which serves as an internal *msh2Δ/msh2Δ* control (Figure 5D) (Dataset S1). Thus, as with *pol3-01/pol3-01 msh6Δ/msh6Δ* diploids, there is strong selection pressure for *pol2-P301R/POL2 msh2Δ/msh2Δ* cells to suppress their mutator phenotype, even though they do not show an obvious initial growth defect.

**Figure 5.**
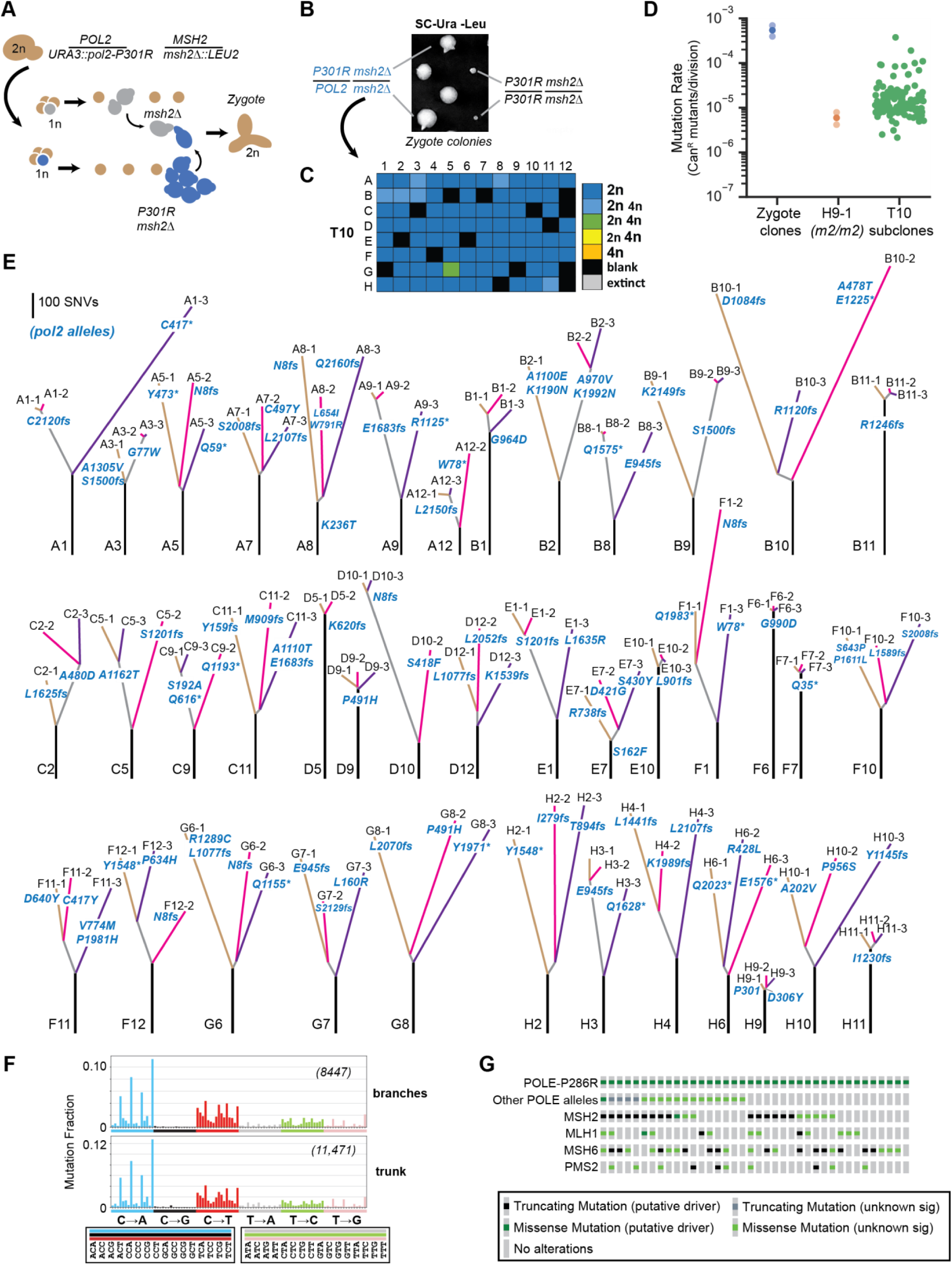
Modeling adaptation of POLE-P286R MMR-deficient cancers to *EEX*. A) Isolation of *pol2-P301R/POL2 msh2Δ/msh2Δ* zygotes. *POL2 msh2Δ* spores slowly divide once on SC-Ura-Leu media, allowing mating with more rapidly dividing *pol2-P301R msh2Δ* cells. B) Representative growth phenotypes of the initial zygote colonies (see Figure S8 for all colonies). C) Ploidy of evolved cultures after ten (T10) serial transfers. Box color indicates the ploidy or mixture of ploidies of each culture as depicted in the key to the right (see Dataset S1 for histograms). D) Mutation rates as determined by a canavanine resistance (Can^R^) fluctuation assay (see Dataset S1 for mutation rates and 95% confidence intervals). Solid circles, mutation rates; transparent circles, 95% confidence intervals (for Zygote Clones only). E) Phylogenetic trees based on whole genome sequencing of the subclones. Line length, the number of shared or unique SNVs; lettering at the bottom of each tree, culture position in 96 well plate; lettering at the end of each branch, subclone ID number; *pol2* mutations are in blue lettering: *fs*, frameshift; *, nonsense mutation. F) Spectra of grouped mutations from trunk and branches. Format is the same as in Figure 4. Trunk spectrum determined from all trunk mutations combined. Branch spectrum determined from combined mutations from branches with truncating mutations at E945 or before (A1-3, A5-1, A5-2, A5-3, A8-1, A12-2, B3-3, C11-1, F1-2, F1-3, F12-2, G6-2, G7-1, H2-2, H2-3, and H3-1/H3-2). G) Oncoprint of POLE-P286R tumors on cBioPortal showing presence of additional *POLE* alleles and mutations affecting MMR components (see Dataset 1).

To understand the basis for the antimutator phenotypes, we sequenced the genomes of 119 subclones from 40 evolved cultures. The subclones averaged 665±178(SD) SNVs and 225±35(SD) Indels, corresponding to 60 SNVs/Mb and 20 Indels/Mb (Dataset S2). We annotated all variants (Dataset S2) and constructed phylogenetic trees using the SNVs (Figure 5E). Secondary mutations in *pol2* were found in all subclones, suggesting that they are adaptive (Figure 5E, Dataset S2). In 6 out of 40 evolved cultures, the *pol2* mutation preceded the divergence of the subclones (B1, B11, D9, F6, F7, H11). As they diverged, the subclones accumulated few additional mutations, providing strong evidence for a direct role of the *pol2* mutations in mutator suppression. Altogether, we observed 88 candidate adaptations in *pol2*, including 45 frameshifts, 23 nonsense mutations, and 36 missense (or combinations of missense) mutations. Two mutant alleles, *I279fs* (H2-2) and *D306Y* (H9-2,3), were close enough to *P301R* to confirm they reside on the same chromosome. The abundance of truncation mutations suggests that inactivation of the *pol2-P301R* allele allows cells to use WT *POL2* as the sole source for Polε, thereby dramatically lowering their mutation rate. This mirrors how the combination of nonsense and antimutator mutations suppresses the mutator phenotype of some *pol3-01/pol3-01 msh6Δ/msh6Δ* cultures. The abundance of truncating *pol2* mutations in these strains also indicates that the observed missense mutations may either inactivate the mutator polymerase or function as antimutators.

The central role of the *pol2* mutations in the evolution of these cultures helps to interpret the structure of the phylogenetic trees. Since most *pol2* mutations occurred after divergence of the subclones, most of the 300±126(SD) shared mutations in the trunks likely arose prior to zygote formation in the *pol2-P301R msh2Δ* haploids. The magnitude of the mutator phenotype of these haploid cells is likely similar to *pol2-P301R/pol2-P301R msh2Δ/msh2Δ* diploid cells, which formed small colonies (Figures 5B, S8). In the mathematical model of diploid *EEX*, homozygous inactivation of essential genes limits growth (Herr *et al.* 2014). A terminal colony size of 1 × 10^5^ cells corresponds to a mutation rate of ∼5 × 10^−3^ Can^R^ mutants per division, which is 10 times greater than the observed *pol2-P301R/POL2 msh2Δ/msh2Δ* mutation rate (5.43 × 10^−4^ Can^R^ mutants/division). We can deduce the genome-wide mutation rate of *pol2-P301R/POL2 msh2Δ/msh2Δ* cells from the minimal internodal distance between branch points in trees that rapidly diverged and acquired independent *pol2* mutations (Dataset S2). The lowest number of shared SNVs between any two subclones was 24. Pairs of subclones from 8 cultures (A5, A7, A8, C11, F10, G6, G8, and H6), with similar low increments of shared SNVs, averaged 30±7 (SD) SNVs/diploid genome. This would place the mutation rate of *pol2-P301R/pol2-P301R msh2Δ/msh2Δ* diploids at ∼300 SNVs/division and the *pol2-P301R msh2Δ* haploids cells at 150 SNVs/division, easily accounting for the observed number of mutations in the trunks of the phylogenetic trees.

If a mutation rate of 30 SNVs/division continued unabated in *pol2-P301R/POL2 msh2Δ/msh2Δ* cells, the subclones would have accumulated ∼3600 mutations over 120 generations. However, the average branch length was 418(±188 SD). A truncating mutation in *pol2-P301R* that nullifies the mutator phenotype should lower the mutation rate to that observed in *msh2Δ/msh2Δ* cells (5 × 10^−6^ Can^R^ mutants/division)(see H9-1, Figure 5C, Dataset S2), which we estimate would be ∼0.3 mutations/diploid genome/division. A hundred cellular generations at this suppressed mutation rate would produce 30 SNVs, equal to the number that occurs in just one division of the original mutator strain. Subtracting this value from the average branch length, suggests that the average adaptative event occurred by ∼ 13 cellular generations, before the colony reached 10^4^ cells (2^13^). Thus, adaptive mutations in *pol2-P301R* occur early during the evolution of these strains, substantially moderating the mutation burden.

Since the bulk of mutations in branches with strong *pol2* null alleles come from these early divisions, we pooled them to determine the mutation spectrum produced by the *pol2-P301R/POL2 msh2Δ/msh2Δ* mutator phenotype (Figure 5F). The spectrum from the branches matches the spectrum of the pooled mutations from the trunks (cosine similarity of 0.994) despite likely differences in the magnitude of their mutator phenotypes (Figure 5F). We also compared the branch mutation spectrum to all mutational signatures from the Catalogue of Somatic Mutations in Cancer (Cosmic) database (Alexandrov *et al.* 2020) to test how well these yeast cells recapitulate the mutational processes occurring in human disease. We found the highest similarity with Cosmic mutational signature 14 (0.793), which results from the synergy between *POLE* and MMR mutator alleles (Haradhvala *et al.* 2018; Alexandrov *et al.* 2020) (Table S1). Thus, our analysis of the phylogenic trees of evolved *pol2-P301R/POL2 msh2Δ/msh2Δ* strains reveal a severe mutator phenotype that resembles the mutagenesis in cancers with combined *POLE* and MMR deficiencies. In this model, we show that adaptation readily occurs through suppression of the *pol2-P301R* allele. In nearly every culture, the adaptation is polyclonal, resulting in a genetically diverse population of cells that survive extinction.

## Discussion

Cell lineages with very high mutation rates face a progressive loss of fitness and even extinction unless they adapt by becoming polyploid or acquiring antimutator mutations. The competition between polyploids and antimutators during the evolution of mutators is influenced by the rate at which they arise and the extent to which they preserve fitness.

In our study of *pol2-4 msh2Δ* haploids, we found that two-thirds of cells sampled from the perimeter of the initial colonies (<20 cellular generations) were inviable, illustrating the strong selection pressure for *eex* mutants. At this early stage, polyploids and antimutators were present with similar frequencies, which is in keeping with the comparable rates of spontaneous polyploidization of haploid yeast (7 × 10^−5^ per division) (Harari *et al.* 2018) and gene inactivation of *pol2-4 msh2Δ* cells (1 × 10^−4^ Can^R^ mutants/division). Diploids frequently emerge in WT haploid evolution studies, with a fitness advantage of around 3.5% per generation (Gerstein *et al.* 2006; Venkataram *et al.* 2016; Fisher *et al.* 2018). The fitness advantage of polyploids during the evolution of haploid mutators is likely much larger and constantly increasing as mutations accumulate, as suggested by studies with chemically mutagenized haploid yeast (Mable and Otto 2001). As the mutator parent strain loses fitness and polyploids and antimutators compete for supremacy, continued mutation accumulation in antimutators may place them at a competitive disadvantage. Even more consequential, the antimutator mutations may inherently compromise fitness by diminishing polymerase processivity. In some cases, antimutators may even cause DNA replication stress and a need for homologous recombination, which would favor the emergence of polyploid variants within the antimutator population.

While spontaneous polyploids and antimutators also arise during the evolution of strong diploid mutators, antimutators dominate nearly every culture. This may reflect stark differences in how frequently the two types of mutants arise. The rate of tetraploidization of diploid yeast, to our knowledge, has not been reported, but our data provide an estimate based on the following logic. Sequencing shows that a single adapted clone rose to prominence in each *pol3-01/pol3-01 msh6Δ/msh6Δ* culture. We observed that 1% of these cultures became tetraploid during their evolution. Mutator suppression of *pol3-01/pol3-01 msh6Δ/msh6Δ* cells requires two independent *pol3* mutations, whereas polyploidization is a singular event. Biallelic attenuation of the *pol3-01* mutator phenotype will occur with a rate no greater than the square of the *per* gene mutation rate, which for *pol3-01/pol3-01 msh6Δ/msh6Δ* cells is ∼(1 x 10^−3^)^2^ or 10^−6^. If we assume that tetraploids and antimutators have a similar stabilizing effect on the loss of fitness, the rate of tetraploid formation may be no greater than 10^−8^. Of course, the rate of tetraploid formation could potentially be higher if tetraploids are inherently less fit than diploid antimutator strains. Growth competition studies between WT tetraploids and diploids suggests this may be true (Gerstein *et al.* 2006). Tetraploids suffer higher rates of genomic instability and depend more heavily on factors involved in homologous recombination (Storchova *et al.* 2006). In the context of a mutator phenotype, a higher rate of homologous recombination in tetraploids may limit buffering of deleterious mutations by promoting loss of heterozygosity. Diploid antimutators are already buffered from mutations and may retain that protection better than tetraploids if they indeed have lower recombination rates.

The ease with which *pol3-01/pol3-01 msh6Δ/msh6Δ* cells adapted to *EEX* despite being homozygous for the mutator alleles, suggests that similar events may be common in cancers with heterozygous *POLE* mutations. We modeled this scenario using *pol2-P301R/POL2 msh2Δ/msh2Δ* cells, which incur 2.7 SNVs/Mb/division (30 SNVs/11Mb). This level of mutagenesis imposes a strong selection for suppressor mutations, and we found a wide variety of suppressor mutations that reduced mutation rate as much as a 100-fold. The average mutation burden of these adapted subclones (60 SNVs/Mb) is less than the clonal mutation burden of many ultra-mutated tumors (>100 SNVs/Mb). Half of *POLE-P286R* tumors curated on cBioPortal (Cerami *et al.* 2012) carry clonal, secondary mutations in *POLE* (Figure 5G, Dataset S1). Intriguingly, four of these carry truncating *POLE* mutations including TCGA-IB-7651 (X1335_splice), whose mutation spectrum reflects synergy between *POLE* and MMR defects (Haradhvala *et al.* 2018). The short-read sequencing used for these samples does not reveal whether the mutations are co-linear with *P286R*. But there are only two possibilities: the mutations either suppress the mutator or WT allele. Yeast cells that only express *pol2-P301R* have a 27-fold higher mutation rate (1.4 × 10^−4^ Can^R^ mutants/division) than cells that are heterozygous for the allele (Kane and Shcherbakova 2014). Thus, truncating *POLE* mutations in the WT allele may enhance the *POLE-P286R* mutator phenotype. The TCGA-IB-7651 tumor carries four other mutations in *POLE* besides the truncating mutation, making it a reasonable candidate for mutator suppression regardless of which *POLE* allele carries the truncating mutation. The mutational signatures of the other three tumors with truncating *POLE* mutations (TCGA-AX-A05Z, P0010967, P0005824) have high mutation loads but lack a mutational signature of combined *POLE* and MMR defects (Haradhvala *et al.* 2018). Thus, these truncating mutations may enhance the *P286R* mutator phenotype. Given their high mutation loads, we predict that these tumors may also experience selection pressure for *P286R* suppressor mutations. However, since *POLE* is essential, adaptations in these types of tumors may be limited to missense mutations that restore Polε fidelity.

The adaptive missense mutations in our study encode amino acid changes that may inactivate the polymerase or attenuate the mutator phenotype. They group together in different locations within the Pol2p structure (Swan *et al.* 2009; Parkash *et al.* 2019), suggesting discrete mechanisms of suppression (see Figure S9 for details). The most striking antimutator candidate is D306Y, which is adjacent to P301R in the exonuclease structure (Figure S9B) and may even be allele specific. Together, the truncating and missense mutations argue that *POLE* cancer cells have numerous ways to escape error-induced extinction. The frequent emergence of independent *eex* mutants in the same evolved culture indicates that polyclonal adaptation may be common in *POLE* tumors, potentially obscuring the critical role of suppressor mutations in preventing error-induced extinction of the cancer and preserving genetic diversity for tumor evolution.

Other changes that attenuate the mutator phenotype may also be possible. In yeast, deletion of *DUN1* suppresses Polδ (Datta *et al.* 2000; Mertz *et al.* 2015) and Polε mutator phenotypes (Williams *et al.* 2015) in haploid MMR-proficient cells. The *dun1Δ* antimutator effect involves a reduction in dNTP pools, since restoration of dNTPs by deletion of *SML1*, a Dun1 target, restores the mutator phenotype (Williams *et al.* 2015). Like intragenic mutations in polymerase genes predicted to compromise DNA binding, mutations that lower dNTP pools may enhance polymerase dissociation from the primer•template, permitting extrinsic proofreading (Williams *et al.* 2015). Although heterozygous variants in *DUN1* are observed in some *pol3-01/pol3-01 msh6Δ/msh6Δ* lineages, they do not explain the observed variation in mutation rates between subclones. While MMR-proficient *pol3-01* cells appear to activate the check-point (Datta *et al.* 2000), this activation may depend, in part, on processing of mismatches by MMR (Reha-Krantz *et al.* 2011). Therefore, in the absence of MMR, the contribution of Dun1p to the *pol3-01* mutator phenotype may be substantially reduced. Recent evolution studies in diploid yeast pointed to unconventional mutator phenotypes caused by heterozygous mutations in house-keeping genes (Coelho *et al.* 2019). The vast phenotypic space created by heterozygous mutations and their epistatic interactions remains largely unexplored. If mutation rate can be enhanced by such mutations, perhaps it may be attenuated as well.

## Materials and Methods

### Culture conditions

Yeast propagation and tetrad dissection followed standard procedures (Sherman 2002). Unless otherwise noted, reagents were purchased from Fisher Scientific or Sigma. For rich liquid media, we used YPD (1% wt/vol yeast extract, 2% wt/vol peptone, 2% wt/vol dextrose). For rich solid media we used YPD or synthetic complete (SC) [6.7 g Difco yeast nitrogen base without amino acids, 2% wt/vol dextrose, 2 g/L SC amino acid Mix (SCM) (Bufferad)] supplemented with 2% wt/vol agar. For drop-out media, SCM powders lacking specified amino acids were either purchased from Bufferad or made to the same specifications (Kaiser 1994). For haploid mutation rates, we used SC plates lacking arginine (SC-Arg) with 60 μg/mL canavanine (Can). For mutation rates with diploid strains we used SC_MSG-Arg [1.7 g Difco yeast nitrogen base without amino acids or ammonium sulfate, 1 g/L monosodium glutamate (for a nitrogen source), 2% wt/vol dextrose, 2 g/L SCM-arginine)] with 60 μg/mL Can and 100 μg/mL nourseothricin (NTC) (Herr *et al.* 2014). We used similar plates lacking histidine and containing NTC (SC_MSG-His+NTC) to select for mating between *CAN1::natMX* and *can1Δ::HIS3* cells (see below).

### Yeast Strains

The diploid parent strain used for these experiments, AH0401, is a BY4743 derivative engineered to be heterozygous at the *CAN1* locus (Herr *et al.* 2014). One allele carries the *natMX* transgene (encoding resistance to NTC) inserted just downstream of the *CAN1* coding sequence (*CAN1::natMX*), while the other *CAN1* allele is deleted and replaced with *HIS3* (*can1Δ::HIS3*). Selection for *can1* mutants in the presence of canavanine and NTC allows us to score *bona fide can1* mutants without the high background of *can1Δ::HIS3/can1Δ::HIS3* mitotic recombinants (Herr *et al.* 2014). The four WT strains used for generating mapping strains (Figure S1) and for the mating-type switching experiments (Figures S3, S4) were dissected from the same AH0401 tetrad: AH4002 (*MAT*α *CAN1::natMX)*, AH4003 (*MAT***a** *CAN1::natMX)*, AH4010 (*MAT*α *can1Δ::HIS3*), and AH4015 (*MAT***a** *can1Δ::HIS3*). AH12721 is a *POL2/URA3::pol2-4 MSH2/msh2Δ::LEU2* strain isolated by mating freshly dissected spores from AH2801 (*POL2/URA3::pol2-4 MSH6/msh6Δ::LEU2*) (Kennedy *et al.* 2015) and AH5610 (*MSH2/msh2Δ::LEU2*). Zygotes were isolated by microdis-section after 8 hours and allowed to form colonies, which were then genotyped. AH5610 was constructed for this study by transforming AH0401 with a *LEU2* PCR product amplified from pRS416 with Msh2U and Msh2D as described (Williams *et al.* 2013). AH2601 is a previously described *POL3/URA3::pol3-01 MSH6/msh6Δ::LEU2* strain (Lee *et al.* 2019) constructed in the AH0401 background. All strains derived from mating between *pol3-01 msh6Δ* spores used in the evolution experiment were designated AH164_NN, where “NN” refers to their coordinate in the 96-well plate. AH11304 is a *URA3::pol2-P301R* transformant of AH5610 in which a duplicated *POL2* promoter flanks the *URA3* transgene upstream of the *POL2* coding sequence. We engineered the strain using a chimeric URA3::*pol2P301R* PCR product amplified in two fragments from pRS416-POL2 (Williams *et al.* 2013) with Phusion polymerase (New England Biolabs) and the following conditions: 98°C, 1min; 25 x (98°C, 10 sec; 51°C, 30 sec; 72°C, 2 min); 72°C 2 min. One fragment was amplified with pol2U (ATGATGAAAGAGCACAT-TCTATCAAGATAACACTCTCAGGGGACAAGTATAGATTGTACTGAGAGTGCAC) and POL2P301R-r (CATTATTTGATCTACGGCGGAATCCcGGAATTTTAAAGGCGGCTTCGTGG; *pol2-P301R* mutation, lower case). A second *POL2* fragment was amplified with POL2P301R (CCACGAAGCCGCCTTTAAAATTCCgGGATTCCGCCGTAGATCAAATAATG) and pol2S7 (ATGTGGATAACTTGGTCTGCG). The two PCR fragments were gel purified, equimolar ratios were combined without primers in a second PCR reaction for ten cycles (98°C, 10 sec; 55°C, 30 sec; 72°C, 5 min), outside primers pol2U and pol2S7 were added, and the reaction was continued for another 10 cycles. The entire *pol2-P301R* gene was initially confirmed by Sanger sequencing and then by whole genome sequencing of the evolved clones.

### Isolation of eex mutants

For *pol3-01 msh6Δ eex* mutants (Figure S1), we plated large numbers of random haploid spores onto media that selected for the *URA3* and *LEU2* transgenes tightly linked to each mutator allele. The *pol3-01 msh6Δ* cells do not form visible colonies unless they acquire an *eex* mutation. The isolation of *pol2-4 msh2Δ eex* mutants required a slightly different approach. During our initial tetrad analysis of the *pol2-4/POL2 msh2Δ/MSH2* strain, we found that *pol2-4 msh2Δ* cells were not synthetically lethal, as they were in our earlier plasmid shuffling system, but grew slowly (Williams *et al.* 2013). To isolate *eex* mutants, we simply re-plated the slow-growing double mutant spore clones for faster-growing colonies. Haploid yeast cells grow vegetatively as one of two mating-types, *MAT***a** and *MAT*α, which can mate to form *MAT***a**/*MAT*α diploids. To isolate mapping strains, we mated each *eex* mutant in parallel with AH4002, AH4003, AH4010, and AH4015 and plated the mutants on SC_MSG-His+NTC. Candidates that only mated with one of the four strains were carried forward for genetic analyses.

### Assay for mating-type switching or same-sex mating

We mixed ∼5 × 10^6^ cells of two strains with the same mating type but opposite *CAN1* alleles (*can1Δ::HIS3* and *CAN1::natMX*) in a microtiter plate with YPD and incubated the cells at 30°C for 8 hours without shaking. *CAN1::natMX/can1Δ::HIS3* diploids were then selected on SC_MSG-His+NTC media. Wild type and *pol2-4 msh6Δ* mutator cells from the T1 transfer were used for Figure S3. Wild-type cells picked from individual colonies were used for Figure S4. Mating type PCRs were performed as described (Harari *et al.* 2018).

### Explaining the similar rates of mating-type switching and same-sex mating

Our finding that diploid colonies occur with similar frequencies indicates that a dsDNA break likely initiates both mating-type switching and same sex mating. In *MAT***a** cells the dsDNA break is repaired by gene conversion from the HML locus, converting the cell to a *MAT*α cell, which then mates with an adjacent *MAT***a** cell to form a *MAT***a***/MAT*α diploid. In *MAT*α cells, the dsDNA break likely disrupts expression of the MATα2 repressor, which normally suppresses the expression of genes required for the *MAT***a** program (Strathern *et al.* 1981). These phenotypic “*MAT***a***”* cells then mate with adjacent *MAT*α cells and repair the break through intra- or inter-chromosomal recombination.

### Evolution Experiments

To obtain *URA3::pol2-4 msh2Δ::LEU2* double mutant spores we dissected tetrads on SC-Ura-Leu. After two days of growth, we isolated single cells from the perimeter of each colony by microdissection and moved them below the colony to form new colonies (Figure S2A). We then suspended the main colonies in 100 μls of H_2_O in 96-well format (Figure 1). Random empty wells were interspersed among the samples to monitor for contamination. We made glycerol stocks from part of the cell suspension. From the remaining cell suspensions, we diluted the cells 1:50 in YPD, and grew 3 hours before removing aliquots for the first flow cytometry measurement after which the cultures were grown to saturation and then subcloned as described below.

To isolate mutator diploids for evolution experiments, AH2601 or AH11304 tetrads were dissected on SC-Ura-Leu plates and spores were separated from each other by ∼100 μm (the width of two diameters of the dissecting needle). After 2 to 3 divisions, dividing cells from different tetrads were placed next to each other to allow mating. Half of the pairings produced zygotes after 2 to 3 additional divisions. Zygotes were moved to a defined coordinate on the plate and allowed to form a colony. In our initial experiment with AH2601, we noted that Ura^-^ Leu^+^ spores germinated and divided once on this media and that we could isolate *pol3-01/POL3 msh6Δ/msh6Δ* heterozygotes by crossing them with more rapidly dividing Ura^+^ Leu^+^ cells. We used this approach to enrich for *pol2-P301R/POL2 msh2Δ/msh2Δ* strains in the AH11304 crosses. We also obtained numerous *pol2-P301R/pol2-P301R msh2Δ/msh2Δ* zygotes in the process, which allowed us to compare the relative colony forming capacity of the two genotypes. Colonies were picked from the plate and suspended in 100 μls of H_2_O. We used 30 μls of the cell suspensions to inoculate 500 μls cultures, 40 μls for glycerol stocks, and the remaining cell suspension to measure mutation rates (see below). For AH11304 derived zygotes, we reserved 10 uls for *pol2P301R* genotyping. The *pol2P301R* allele creates a sequence (…AATTCCgGG…) that is one bp from a DraI site (CCCGGG). To genotype *pol2-P301R* we amplified *POL2* with pol2-P301R-dctF (TGTGGTAATGGCATTTGATATAGAAACCACGAA-GCCGCCTTTAAAATcCC), which has a mismatch (lower case) that creates a DraI site, and pol2-P301R-dctF (GGATACTCCGGTTTCGGTGTATACTCAAAGTCTTCAATATCC-TCAGAGA). We digested the amplified products with DraI in the NEB Cutsmart buffer and resolved the fragments on a 3% agarose gel. The 180bp product is cleaved into fragments of 50 and 130bp if the *pol2-P301R* allele is present.

In both evolution experiments, the strains were propagated at 30°C with constant shaking in sterile Nunc 96 deep-well storage blocks (Thermo Scientific, 260252), covered with an AeraSeal (Excel Scientific) gas-permeable disposable plate sealer. Each well had 500 μls of YPD and a single 3 mm glass bead to maintain cells in suspension. Sub-culturing steps were performed by transferring 1 ul volumes from saturated cultures to 500 μls of fresh YPD using an 8-channel pipettor. Frozen -80°C glycerol stocks of the cultures were made every three transfers in 96-well format. To recover cultures from storage, the plates were first thawed at room temperature and then 5 μls were transferred into 250 μl of fresh media.

### Mutation rates

Mutation rates by fluctuation analysis were performed essentially as described (Herr *et al.* 2014). To determine the mutation rate of the zygote colonies, 10 μls of the original suspension (see above) were spot-plated directly on SC_MSG-Arg+CAN+NTC plates. Another 10 μls were used for a series of 10-fold serial dilutions. With each dilution, 10 μls were plated on the mutation rate plate and 10 μls on SC media to estimate the total number cells in each colony (Nt). Only half of the zygote colonies had the *CAN1::natMX/can1Δ::HIS3* geno-type suitable for mutation rate measurements. Thirty colonies with similar Nt values were used as replica colonies to estimate the average number of mutational events (*m*) by maximum likelihood using the newton.LD.plating function in the R package rSalvador (Zheng 2015). The confint.LD.plating function was used to calculate confidence intervals. For mutation rates of all subclonal isolates (Figure 4A, S2), replica colonies were obtained by plating serial dilutions onto SC media. For mutation rate calculations, we typically used mutation counts from 8 replica colonies. In some assays of T1 subclones in Figure 4A, 1 or 2 replica colonies had very low mutation counts (0 or 1 mutants vs 100’s). These clear outliers were censored under the assumption that they represented an antimutator or polyploid subclone. Cohorts with more than 2 such outliers were censored completely.

### Flow Cytometry

Overnight cultures were diluted 1:50 in fresh YPD and grown for 3-5 hours. 100 μls of cells were recovered by centrifugation and washed twice in an equal volume of water by resuspending them and centrifuging again. We then resuspended the cells in 70% ethanol and incubated them at least 1 hour at room temperature or overnight at 4°C. We recovered the cells again by centrifugation, washed once with water, and then resuspended them in 50 mM sodium citrate with RNaseA (10 μg/mL). The cells were heated to 95°C for 15 minutes and then incubated for 3 to 4 hours at 37°C. The cells were recovered by centrifugation and resuspended at a concentration of 10^6^ cells/mL in 50 mM sodium citrate with 2 μM Sytox Green (Life Technologies). After 1 hour at room temperature, the cells were sonicated for 30 seconds with 1 second pulses in a Misonix cup horn sonicator and then analyzed on a BD Accuri c6 flow cytometer using the FL1 detector. Plots in Dataset S1 are organized in the 96 well-format used for each experiment. Histogram plots of the data were generated using the c6 software, which allows multiple plots to be overlaid. Plots used as controls are indicated at the top of each experiment.

### Genome Sequencing and Analysis

We performed whole genome sequencing of sub-clones from the AH164 evolved cultures as described (Herr *et al.* 2014; Lee *et al.* 2019). We used a custom sequencing analysis pipeline (eex_yeast_pileline.sh) that operates in a Unix command-line. It aligns the sequencing reads against a repeat-masked version of the R64-1-1 assembly of the S288C *S. cerevisiae* genome using the Burrows-Wheeler Aligner (0.7.17). Discordant and split reads were removed using Samblaster (0.1.24). Picard tools (2.21.9) Ad-dorReplaceReadGroups was used to add information to the header needed for later steps. After indexing the BAM files with Samtools (1.8), we sequentially processed them with the Genome Analysis Toolkit (GATK3) RealignerTargetCreator, IndelRealigner, LeftAlignIndels, BaseRecalibrator, and PrintReads to minimize false variant calls. We used SamTools to make a pileup file and then VarScan (v2.3.9) mpileup2snp and mpileup2indel to call SNPs and indels, respectively. We used a variant frequency cutoff of 0.1 for SNPs to demonstrate that the sequenced genomes were indeed from 2n and not 4n cells (Dataset 2). We used a variant frequency cutoff of 0.3 for indels to minimize false positives at repetitive sequences due to frameshift errors during library preparation and filtered out SNPs and indels that were present in the parental strains. For AH11304 evolved cultures we prepared DNA sequencing libraries with the NEBNext Ultra II FS DNA library kit, which utilizes enzymatic fragmentation instead of sonication. The same sequencing analysis pipeline was used except that during the variant calls, we used a variant frequency cut-off of 0.22 for SNPs and 0.3 for indels.

We wrote a python script to compare variants found in subclones from the same culture and create lists of shared and unique mutations found in each strain. The program only assesses positions in the genome with 18-fold depth coverage in all subclones (Dataset S2). We annotated the SNP lists using Annotate 0.1 (Pashkova *et al.* 2013) and the Indel lists with a modified version of the program that reports the 3′ peptide produced by the frameshift (anno-tatefs2.py). We used the SNP lists to determine the 96-trinucleoide mutation spectra using snv-spectrum (https://github.com/aroth85/snv-spectrum). We wrote a python script which creates a 2 x 96 table of the output of snv-spectrum for two different variant lists and then estimates *p*-values of the hypergeometric test (Adams and Skopek 1987) and the cosine similarity test, under the null hypothesis that they came from the same population.

Cosine similarity, commonly used in mutation signature analyses, is a measure of the angle between two vectors projected into an n-dimensional space, where n is the number of categories used for the comparison. In this case, the categories are the 96 trinucleotide contexts. The number of mutations at each context together define the direction of the vector. The closer two spectra, the smaller the angle, and the closer the cosine similarity is to 1. The statistical significance of such comparisons is typically not reported. However, the hypergeometric test, which is an expansion of Fisher’s exact test, has previously been used to test the null hypothesis that two spectra are the same. Formally, the hypergeometric probability of each possible contingency table is calculated and the number of tables with a higher probability than the observed table provides a measure of significance. With a 2 × 96 contingency table, an exact calculation of significance becomes computationally impractical but can be estimated using a Monte Carlo approach. Implementation of the test involves construction of 10,000 random 2 × 96 tables that maintain the same row and column totals as the original table (Agresti *et al.* 1979) and then calculation of the hypergeometric probability or cosine similarity of each table (Dataset S2). The fraction of random tables with a hypergeometric probability higher than the original table is an estimate for the probability that two spectra are the same. Likewise, the cosine similarity *p*-value represents the number of random tables whose cosine similarity was higher than the observed cosine similarity. We used the conservative Bonferroni multiple testing correction to estimate the *p*-value cut-off for significance for each set of comparisons (Dataset S2). Finally, we used the SNP lists described above to determine the dN/dS ratios of individual genes and of the entire genome using the R-package *dNdscv* (Martincorena *et al.* 2017) https://github.com/im3sanger/dndscv.

## Acknowledgements

We thank Dr. Maitreya Dunham for use of her flow cytometer and horn sonicator. We thank Dr. Iñigo Martincorena for help implementing *dNdscv*. We thank Drs. Dunham and Scott Kennedy for critically reading the manuscript. We acknowledge funding (R01GM118854) from the National Institute of General Medical Sciences (NIGMS). MBL was supported by the Howard Hughes Medical Institute (HHMI) Gilliam Fellowship for Advanced Study, the National Institutes of Health (NIH) Cellular and Molecular Biology training grant (NIH T32GM7270-39), and the University of Washington Graduate Opportunities and Minority Achievement Program (UW GO-MAP) Bank of America Fellowship. ITD was supported by the UW Molecular Medicine Training Grant (NIH/NIGMS T32GM095421) and Genetic Approaches to Aging Training Grant (NIH/NIA T32AG000057). The content is solely the responsibility of the authors and does not necessarily represent the official views of the NIH, NIA, NIGMS or HHMI.

## Data Availability

Strains and plasmids are available upon request. All whole genome sequencing data is available from the Sequence Read Archive (BioProject ID: PRJNA629499). Supplemental files are available at FigShare. Dataset 1 contains flow cytometry results, mutation rates, and modeling of diploid mutator evolution. Dataset 2 describes whole genome sequencing results of diploid mutator strains. Datasets 1 and 2 can also be found at github (https://github.com/mutatorUW/MutatorEvolution) along with all computer scripts used in this work.

## Supplementary Information

**Table S1:** Similarity of the *pol2/POL2 msh2Δ/msh2Δ* mutation spectra to mutation signatures in cancer.

|  |  |
| --- | --- |
| Signature | Cosine Similarity |
| SBS1 | 0.138 |
| SBS2 | 0.178 |
| SBS3 | 0.485 |
| SBS4 | 0.483 |
| SBS5 | 0.537 |
| SBS6 | 0.239 |
| SBS7a | 0.191 |
| SBS7b | 0.209 |
| SBS7c | 0.175 |
| SBS7d | 0.218 |
| SBS8 | 0.519 |
| SBS9 | 0.456 |
| SBS10a | 0.634 |
| SBS10b | 0.149 |
| SBS11 | 0.344 |
| SBS12 | 0.233 |
| SBS13 | 0.070 |
| SBS14 | 0.793 |
| SBS15 | 0.217 |
| SBS16 | 0.276 |
| SBS17a | 0.053 |
| SBS17b | 0.092 |
| SBS18 | 0.626 |
| SBS19 | 0.301 |
| SBS20 | 0.536 |
| SBS21 | 0.238 |
| SBS22 | 0.046 |
| SBS23 | 0.299 |
| SBS24 | 0.333 |
| SBS25 | 0.447 |
| SBS26 | 0.236 |
| SBS27 | 0.140 |
| SBS28 | 0.159 |
| SBS29 | 0.472 |
| SBS30 | 0.340 |
| SBS31 | 0.258 |
| SBS32 | 0.501 |
| SBS33 | 0.149 |
| SBS34 | 0.075 |
| SBS35 | 0.424 |
| SBS36 | 0.701 |
| SBS37 | 0.276 |
| SBS38 | 0.317 |
| SBS39 | 0.241 |
| SBS40 | 0.623 |
| SBS41 | 0.304 |
| SBS42 | 0.453 |
| SBS43 | 0.052 |
| SBS44 | 0.561 |
| SBS45 | 0.396 |
| SBS46 | 0.266 |
| SBS47 | 0.201 |
| SBS48 | 0.026 |
| SBS49 | 0.071 |
| SBS50 | 0.282 |
| SBS51 | 0.212 |
| SBS52 | 0.180 |
| SBS53 | 0.128 |
| SBS54 | 0.144 |
| SBS55 | 0.082 |
| SBS56 | 0.653 |
| SBS57 | 0.370 |
| SBS58 | 0.375 |
| SBS59 | 0.322 |
| SBS60 | 0.025 |

**Figure S1.**
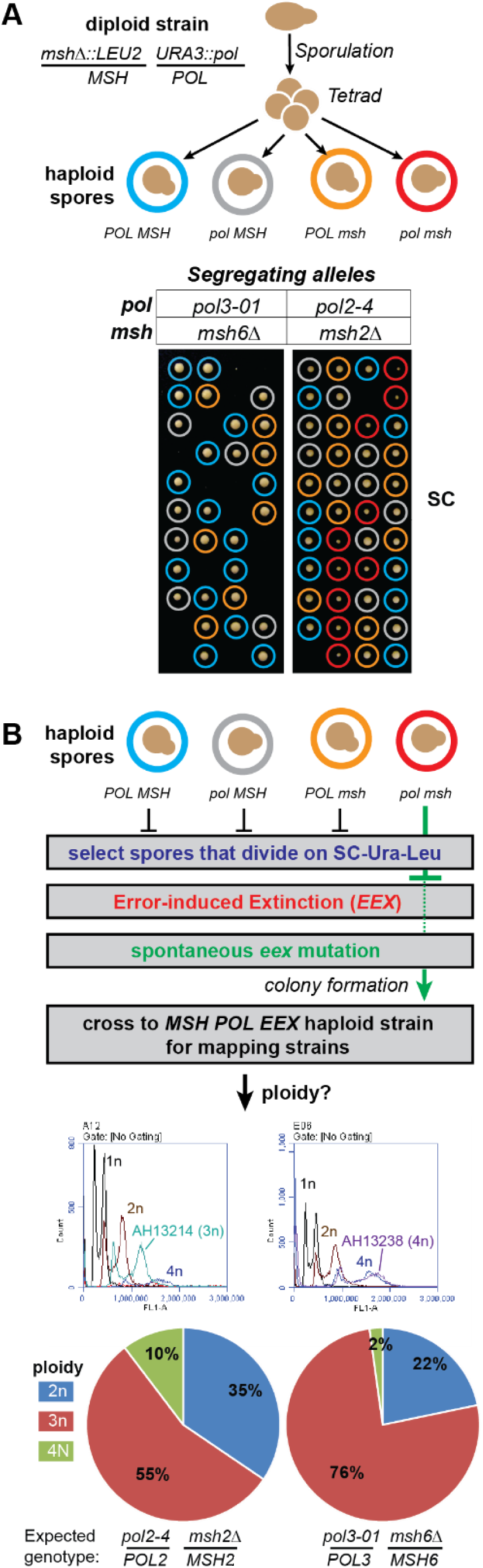
Discovery of spontaneous polyploid *eex* mutants. A) Synthetic growth phenotypes of freshly dissected *pol3-01 msh6Δ* and *pol2-4 msh2Δ* spore clones. *Top*, Schematic of strain genotypes and tetrad dissections. *Bottom*, Images of tetrad dissections after 48 hours of growth. Colored circles indicate genotypes. B) Isolation of *eex* mutants and evidence for polyploidization. *Top*, Scheme for isolating *eex* mutants and the generation of mapping strains. *Bottom*, Flow cytometry traces of two representative polyploid strains (AH13214 and AH13238; see Dataset 1) and 1n, 2n, and 4n controls as well as tally of the ploidies of the mapping strains.

**Figure S2.**
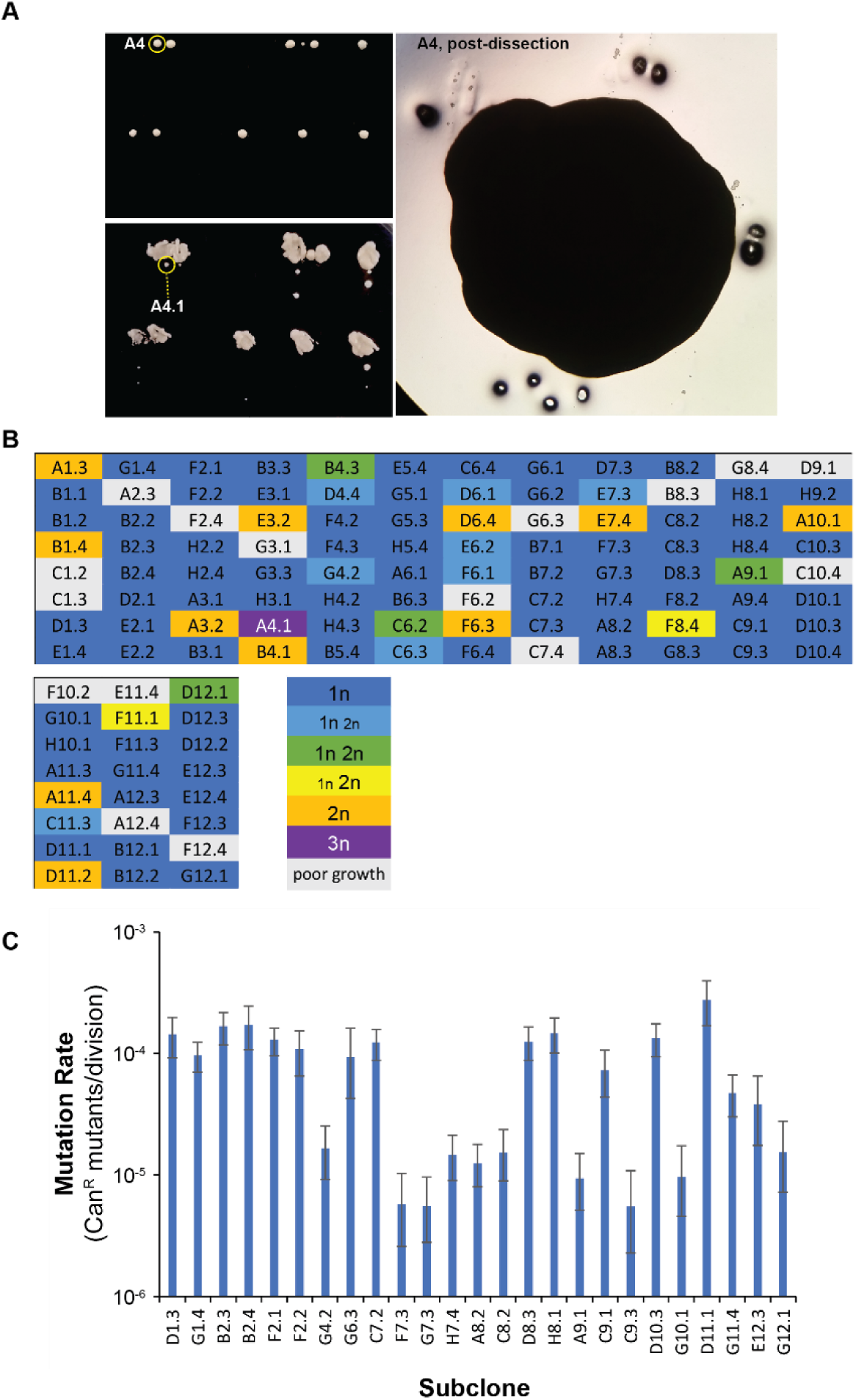
Polyploids and antimutator subclones emerge with similar frequencies during the evolution of *pol2-4 msh2Δ* cultures. A) Representative images of colonies. *Clock-wise from the top left:* initial colonies after 24 hours, A4 (a mixture of 1n and 3n cells) is circled in yellow; close-up of A4 just after removing single cells from the perimeter (holes were punched into the agar to mark origin of cells 1 through 4); colony forming capacity of single cells (A4.1 is circled in yellow). B) Ploidy of colonies derived from single-cell clones (see Dataset S1 for histograms), named for the coordinate of the parent strain in the evolved 96-well plate and the isolate number. C) Mutation rate measurements of haploid cultures as determined by fluctuation analysis of canavanine resistance (Can^R^). Error bars represent 95% confidence intervals. (See Dataset S1 for Table of values.)

**Figure S3.**
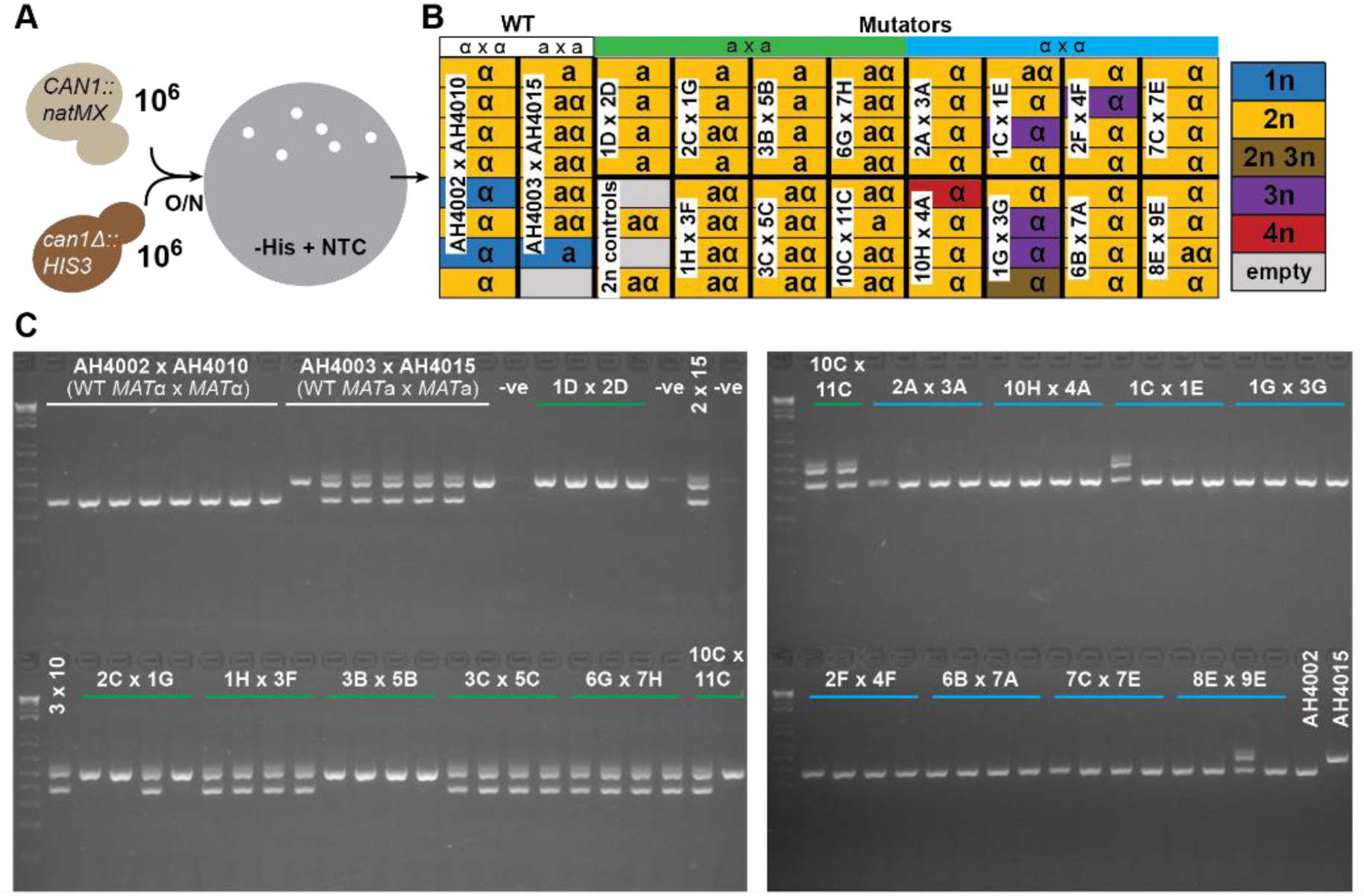
Test for mating-type switching and same-sex mating between mutator cells. A) Experimental design. Same sex haploid cultures with opposite *CAN1* alleles (*CAN1::natMX, can1Δ::HIS3)* were mixed and plated for *CAN1::natMX/can1Δ::HIS3* diploids. B) Summary of flow cytometry and mating-type PCR assays of colonies resulting from the crosses indicated in the white boxes. The mutator cultures used for each cross are indicated by their 96-well coordinate (see Figure 1). The color of the box indicates ploidy. Blue 1n boxes are haploids included as controls (see Dataset S1 for individual histograms). Results of Mating type PCR: **α**, *MATα*; **a**, *MATa* C) Mating-type PCR gels.

**Figure S4.**
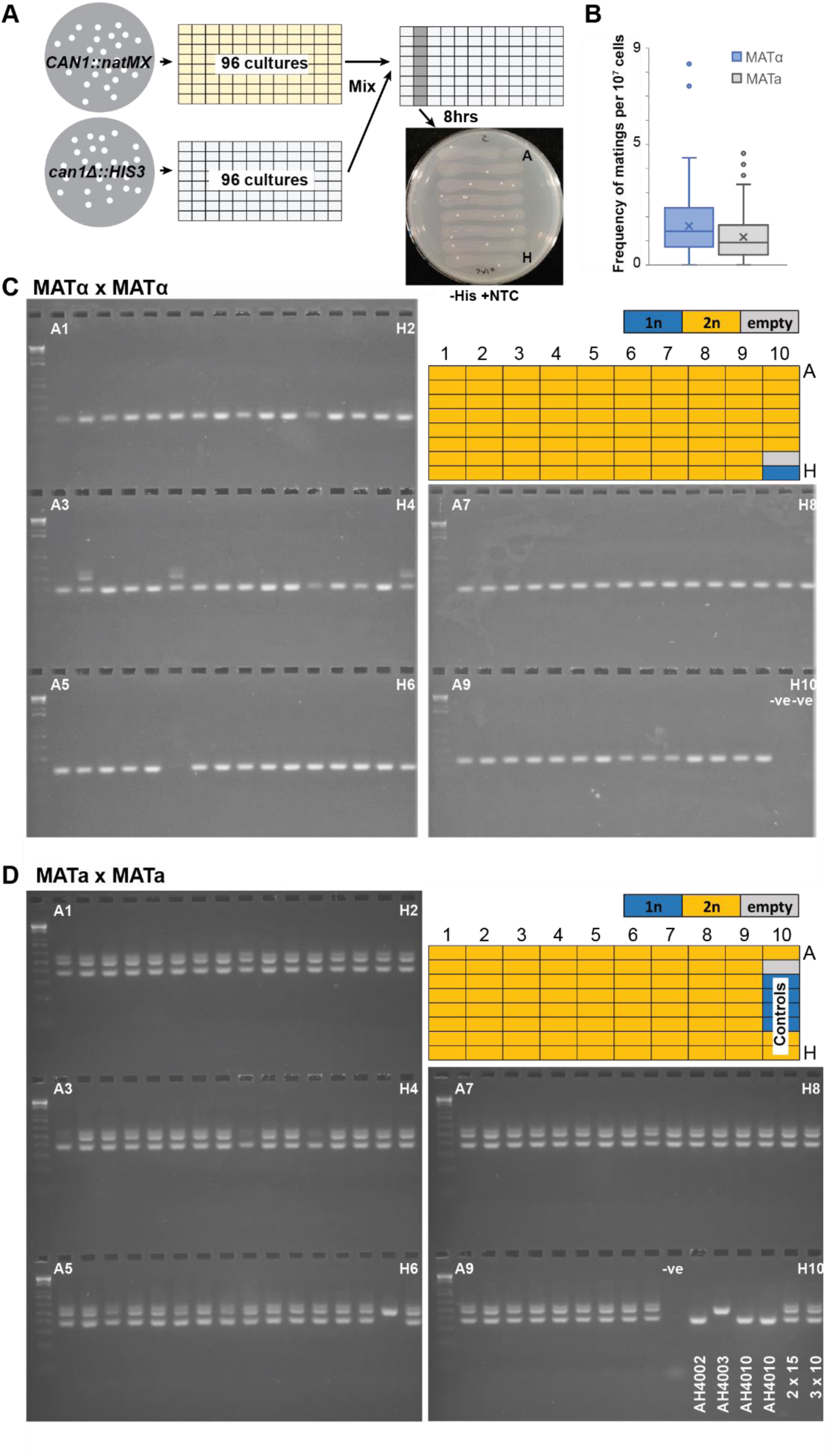
Frequency of mating-type switching and same-sex mating between wild-type cells. A) Experimental design. Same sex haploid colonies with opposite *CAN1* alleles (*CAN1::natMX, can1Δ::HIS3)* were mixed, incubated in media, and plated for *CAN1::natMX/can1Δ::HIS3* diploids. B) Frequencies of His^+^ NTC^R^ colonies per 10^7^ cells. C) Summary of flow cytometry and mating-type PCR assays of colonies resulting from MATα x MATα matings. The color of the box indicates ploidy (see Dataset S1 for individual histograms). D) Summary of flow cytometry and mating-type PCR assays of colonies resulting from MAT**a** × MAT**a** matings. AH4002, AH4003, AH4010 represent haploid strains; 2 × 15 and 3 × 10 are MAT**a**/α control diploids derived from matings between AH4002 × AH4015 and AH4003 × AH4010.

**Figure S5.**
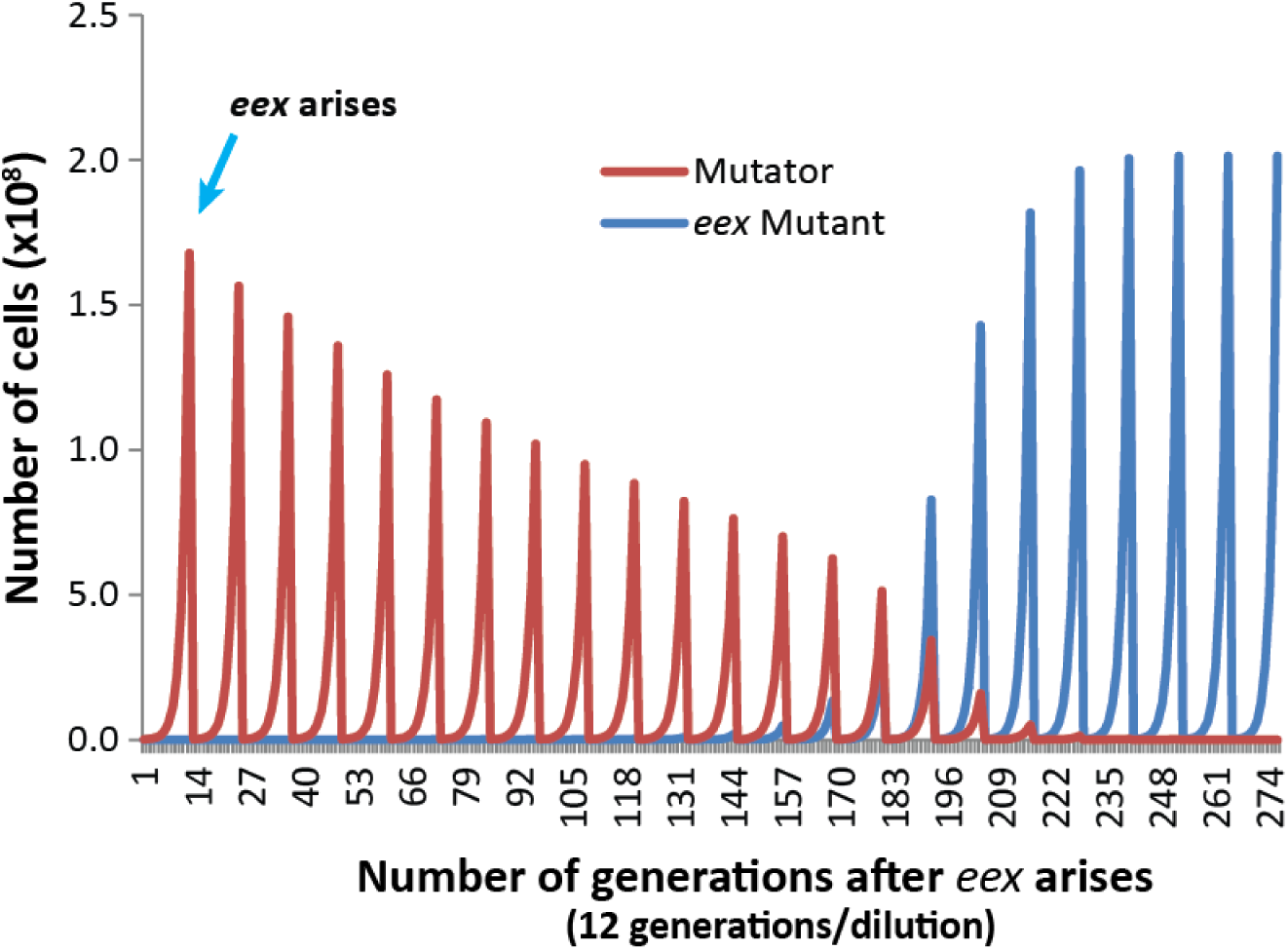
Simulation of *eex* adaption during diploid mutator evolution. Exponential growth of *pol3-01/pol3-01 msh6Δ/msh6Δ* cells (Mutator, red line) and an *eex* mutant (blue line) with a 10-fold lower mutation rate were modeled assuming the only limiting factor for growth was homozygous inactivation of essential genes (see Dataset S1 for calculations). Each peak models a 1:500 dilution of the culture from ∼10^8^ cells to ∼10^5^ cells.

**Figure S6.**
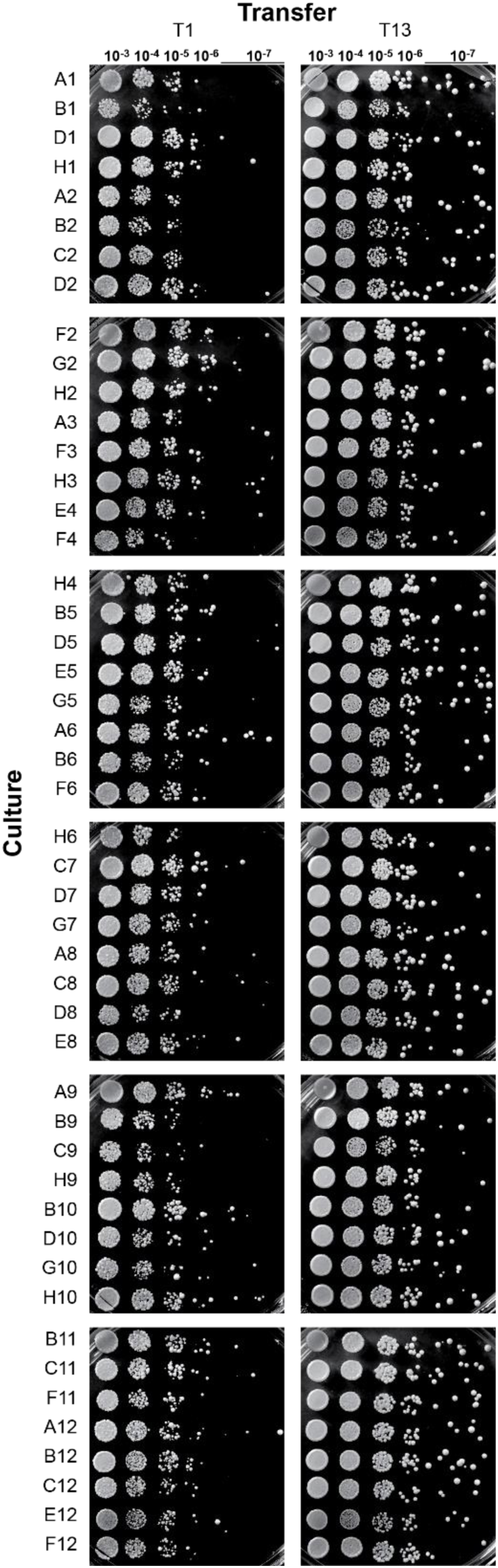
Colony forming capacity of evolved *pol3-01/pol3-01 msh6Δ/msh6Δ* cultures. Ten-fold serial dilutions from overnight cultures were plated on SC plates and incubated for two days at 30°C. T1 and T13 refer to first and thirteenth sub-culturing steps after the initial inoculations. The culture designation on the left refers to the position of the culture in the 96 well plate (see Figure 3).

**Figure S7.**
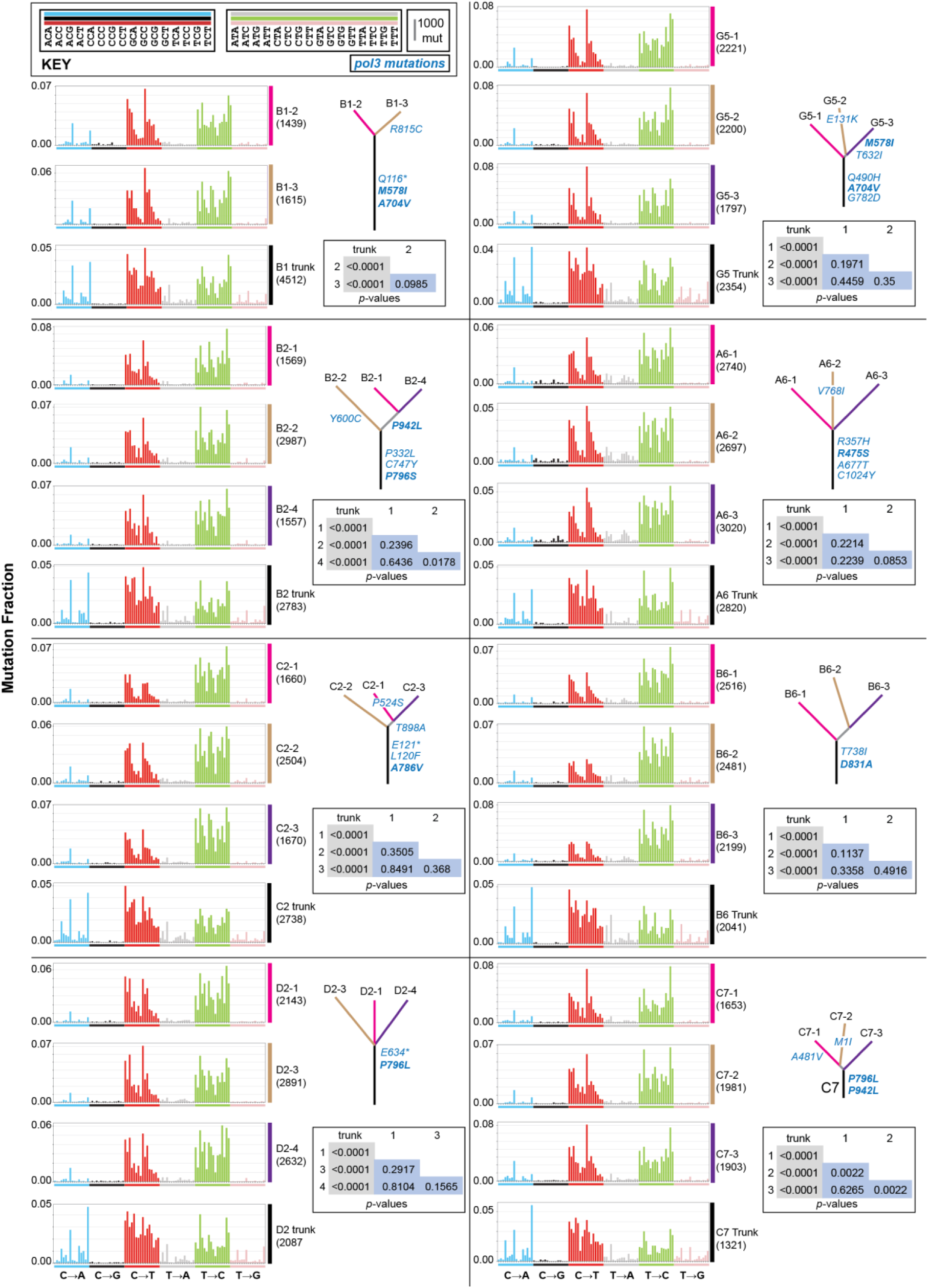
Mutation spectra and phylogenetic relationships of evolved *pol3-01/pol3-01 msh6Δ/msh6Δ* T25 subclones. Mutational variants identified by whole genome sequencing of subclones isolated from different evolved cultures were classified as shared (trunk) and unique. The nucleotide context 5′ and 3′ of each mutational site (represented as the pyrimidine base T or C) were used to classify the 6 mutation types (C:G→A:T, C:G→G:C, C:G→T:A, T:A→A:T, T:A→C:G, T:A→G:C) into 96 trinucleotide contexts. Each mutation type family is color-coded (see Key for color and context order). The bar height for mutation type represents the fraction of the total mutations (to the right of each plot). Phylogenetic trees: line length corresponds to the number of mutations, line color corresponds to mutation spectra, observed *pol3* mutations are in blue lettering (bold represents known or likely antimutators). Inset tables: *p*-values determined by the hypergeometric test.

**Figure S8.**
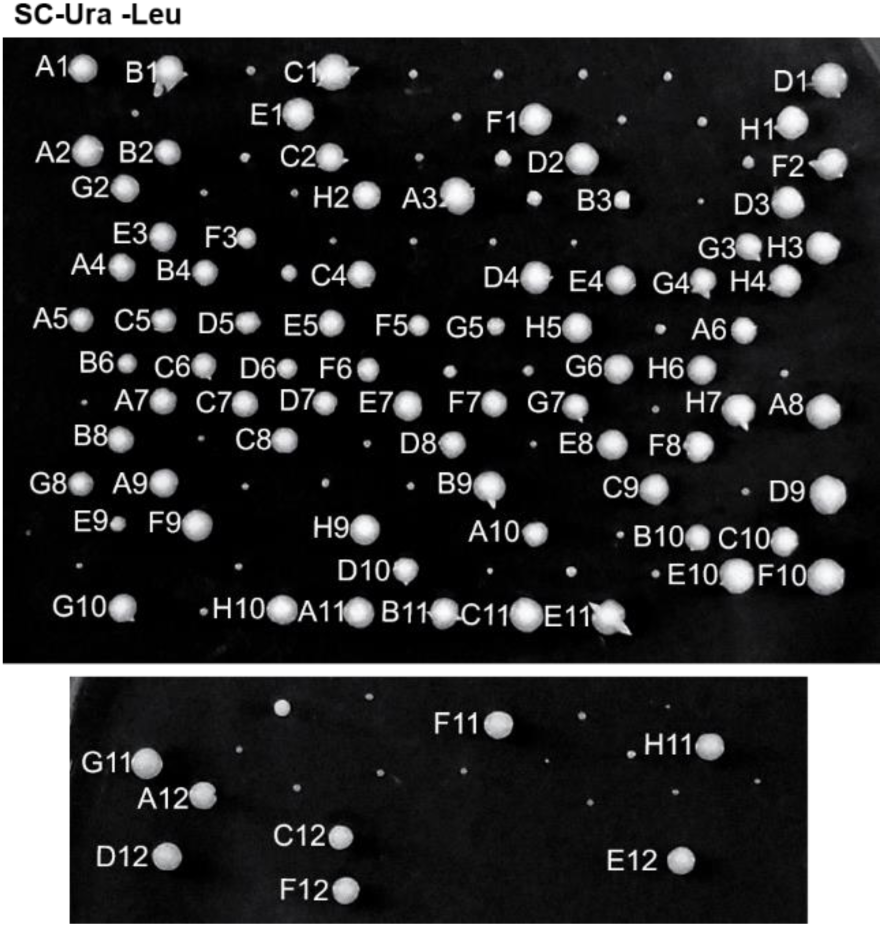
Growth phenotypes of *pol2-P301R/POL2 msh2Δ/msh2Δ* and *pol2-P301R/pol2-P301R msh2Δ/msh2Δ* zygotes. Zygotes colonies were isolated, as described in Figure 5, on synthetic complete media lacking uracil and leucine (SC-Ura-Leu) by mating cells dissected from tetrads of AH11304 (*URA3::pol2-P301R/POL2 msh2Δ::LEU2/MSH2*). Each colony used for the evolution experiment in Figure 5 is labeled to the left by the 96-well coordinate position. Small, unlabeled colonies are derived from *pol2-P301R/pol2-P301R msh2Δ/msh2Δ* zygotes and were not evolved.

**Figure S9.**
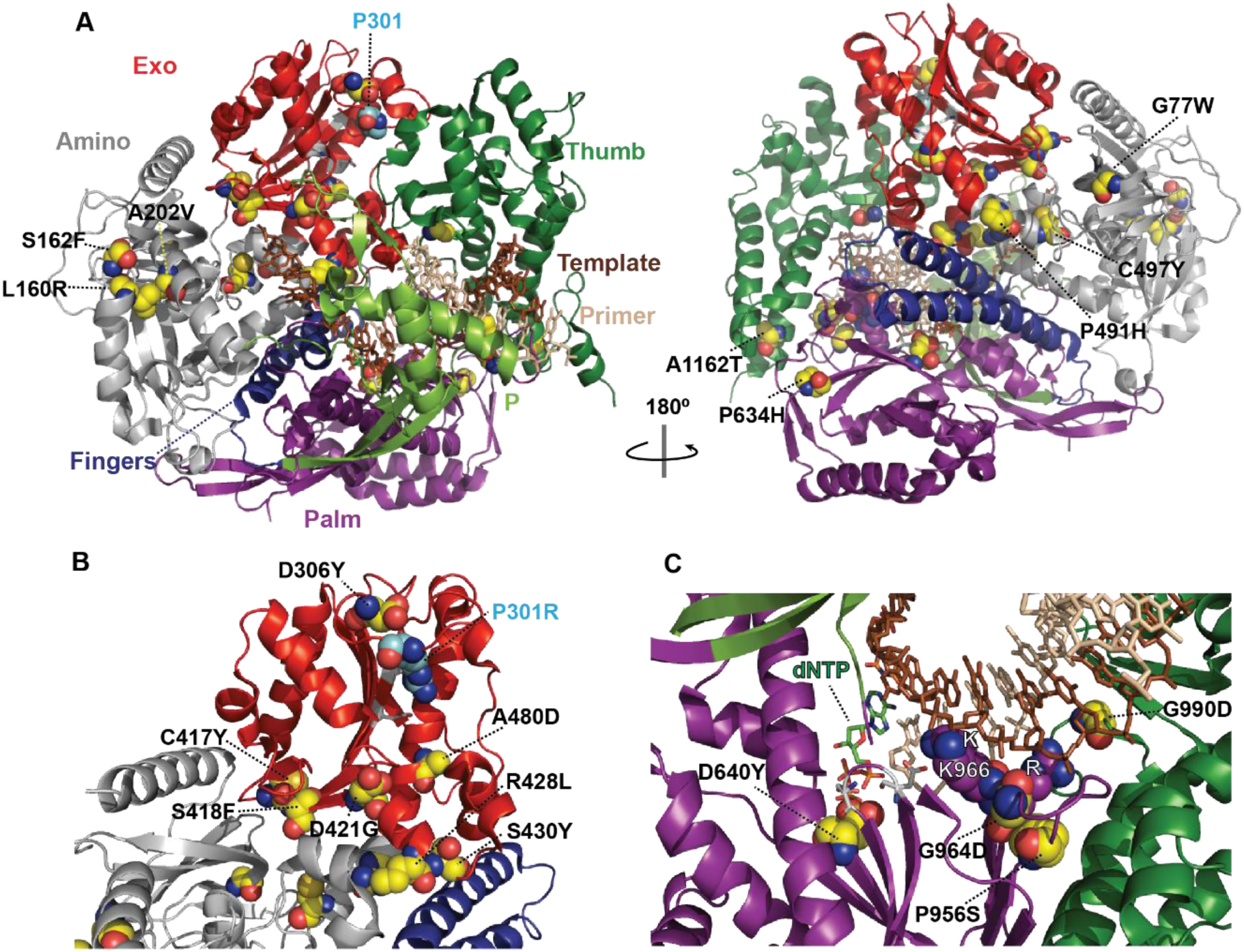
Locations of amino acid substitutions encoded by *pol2-P301R* mutator suppressors. A) Two orientations of the wild-type *S. cerevisiae* Polε structure (Protein database accession code 4M80 (Swan *et al.* 2009)) are shown as ribbon diagrams of the α-carbon backbone (rendered in PyMol). Structural domains are color coded as follows: Amino (gray), Exo (red), processivity (P) (chartreuse), Palm (purple), Fingers (blue) and Thumb (green). Yellow spheres correspond to amino acid substitutions encoded by mutator suppressors. The incoming dNTP is denoted by green CPK sticks. The primer DNA is represented by tan sticks and the template DNA, by brown sticks. Active-site carboxylate side chains are gray CPK sticks coming out of the purple (palm) or red (exo) ribbons. L160R, S162F, and A202V affect interacting residues that play a structural role in the amino terminal domain in a region not previously implicated in fidelity. B) Locations of *eex* substitutions within the Exo domain of the Pol2p-P301R mutant polymerase (Protein database accession code 6i8a (Parkash *et al.* 2019)), which may be candidates for enhancing proofreading function. C) Substitutions positioned to influence dNTP binding and the KKRYA Motif. D640 is 3.21 Å and 3.46 Å from the γ and α-phosphates of the incoming dNTP, respectively. K966, K967(K), and R968(R) from the KKRYA motif, which monitors the minor groove for non-Watson-Crick base-pairing, are depicted as purple CPK space-filling spheres. We previously isolated an antimutator allele affecting K966 (*pol2-K966Q*) (Williams *et al.* 2013). P956S is 3.23 Å from the primary amino group of R968.

**Dataset S1.** Flow cytometry, mutation rates, and modeling of diploid evolution.

|  |  |
| --- | --- |
| Page | Description |
| 1 | Ploidy measurements of *eex* mapping strains described in S1B Fig. |
| 2 | Ploidy measurements of *pol2-4 msh6Δ* evolved cultures described in Fig 1B. |
| 3 | Ploidy measurements of *pol2-4 msh6Δ* T8 subclones described in Fig 1C. |
| 4 | Ploidy measurements of *pol2-4 msh6Δ* subclones described in S2A Fig. |
| 5 | Mutation rate measurements of *pol2-4 msh6Δ* subclones described in S2B Fig. |
| 6 | Ploidy measurements of strains isolated by same sex mating between T1 *pol2-4 msh6Δ* cultures described in S3 Fig. |
| 7 | Ploidy measurements of competition experiment between *pol2-4 msh6Δ* derived diploid and tetraploid strains described in Fig 2B. |
| 8 | Simulation of evolution of *pol3-01/pol3-01 msh6Δ/msh6Δ* evolved cultures described in S4 Fig. |
| 9 | Ploidy measurements of *pol3-01/pol3-01 msh6Δ/msh6Δ* evolved cultures described in Fig 3B. |
| 10 | Ploidy measurements of T25 *pol3-01/pol3-01 msh6Δ/msh6Δ* subclones analyzed in Fig 4 |
| 11 | Mutation rate measurements of T25 *pol3-01/pol3-01 msh6Δ/msh6Δ* subclones described in Fig 4. |
| 12 | Ploidy measurements of *pol2-P301R/POL2 msh2Δ/msh2Δ* evolved cultures described in Fig.5C. |
| 13 | Mutation rate measurements of T10 *pol2-P301R/POL2 msh2Δ/msh2Δ* subclones described in Fig. 5D and *pol2* changes in Fig.5E. |
| 14 | POLE-P286R cancers on cBioPortal depicted in Fig. 5G. |

**Dataset S2.** Whole genome sequencing.

| Dataset S2. Whole genome sequencing. |  |
| --- | --- |
| Page | Description |
| 1 | AH164_B1_SNPs: Sequencing coverage, evolutionary relationships, distribution of variant allele frequencies (VAF), and annotation of SNPs for AH164_B1 *pol3-01/pol3-01 msh6Δ/msh6Δ* subclones. |
| 2 | AH164_B1_Indels: Sequencing coverage, evolutionary relationships, and annotation of Indels for AH164_B1 *pol3-01/pol3-01 msh6Δ/msh6Δ* subclones. |
| 3 | AH164_B2_SNPs: Sequencing coverage, evolutionary relationships, distribution of VAFs, and annotation of SNPs for AH164_B2 *pol3-01/pol3-01 msh6Δ/msh6Δ* subclones. |
| 4 | AH164_B2_Indels: Sequencing coverage, evolutionary relationships, and annotation of Indels for AH164_B2 *pol3-01/pol3-01 msh6Δ/msh6Δ* subclones. |
| 5 | AH164_C2_SNP: Sequencing coverage, evolutionary relationships, distribution of VAFs, and annotation of SNPs for AH164_C2 *pol3-01/pol3-01 msh6Δ/msh6Δ* subclones. |
| 6 | AH164_C2_Indels: Sequencing coverage, evolutionary relationships, and annotation of Indels for AH164_C2 *pol3-01/pol3-01 msh6Δ/msh6Δ* subclones. |
| 7 | AH164_D2_SNP: Sequencing coverage, evolutionary relationships, distribution of VAFs, and annotation of SNPs for AH164_D2 *pol3-01/pol3-01 msh6Δ/msh6Δ* subclones. |
| 8 | AH164_D2_Indels: Sequencing coverage, evolutionary relationships, and annotation of Indels for AH164_D2 *pol3-01/pol3-01 msh6Δ/msh6Δ* subclones. |
| 9 | AH164_G5_SNP: Sequencing coverage, evolutionary relationships, distribution of VAFs, and annotation of SNPs for AH164_G5 *pol3-01/pol3-01 msh6Δ/msh6Δ* subclones. |
| 10 | AH164_G5_Indels: Sequencing coverage, evolutionary relationships, and annotation of Indels for AH164_G5 *pol3-01/pol3-01 msh6Δ/msh6Δ* subclones. |
| 11 | AH164_A6_SNP: Sequencing coverage, evolutionary relationships, distribution of VAFs, and annotation of SNPs for AH164_A6 *pol3-01/pol3-01 msh6Δ/msh6Δ* subclones. |
| 12 | AH164_A6_Indels: Sequencing coverage, evolutionary relationships, and annotation of Indels for AH164_A6 *pol3-01/pol3-01 msh6Δ/msh6Δ* subclones. |
| 13 | AH164_B6_SNP: Sequencing coverage, evolutionary relationships, distribution of VAFs, and annotation of SNPs for AH164_B6 *pol3-01/pol3-01 msh6Δ/msh6Δ* subclones. |
| 14 | AH164_B6_Indels: Sequencing coverage, evolutionary relationships, and annotation of Indels for AH164_B6 *pol3-01/pol3-01 msh6Δ/msh6Δ* subclones. |
| 15 | AH164_C7_SNP: Sequencing coverage, evolutionary relationships, distribution of VAFs, and annotation of SNPs for AH164_C7 *pol3-01/pol3-01 msh6Δ/msh6Δ* subclones. |
| 16 | AH164_C7_Indels: Sequencing coverage, evolutionary relationships, and annotation of Indels for AH164_C7 *pol3-01/pol3-01 msh6Δ/msh6Δ* subclones. |
| 17 | Table of cosine similarity and hypergeometric tests of similarity between *pol3-01/pol3-01 msh6Δ/msh6Δ* mutation spectra. |
| 18 | Output of dNdScv analysis of AH164 evolved cultures reported in Figure 4E. |
| 19 | Evolutionary relationships, variant counts, and annotated *pol2* variants for *pol2-P301R/POL2 msh2Δ/msh2Δ* evolved strains used for Figure 5E. |
| 20 | Annotations of all variants in *pol2-P301R/POL2 msh2Δ/msh2Δ* evolved strains described in Figure 5. |
| 21 | Table of cosine similarity and hypergeometric tests of similarity between *pol2-P301R/POL2 msh2Δ/msh2Δ* mutation spectra. |

